# Signatures of electrical stimulation driven network interactions in the human limbic system

**DOI:** 10.1101/2022.11.23.517746

**Authors:** Gabriela Ojeda Valencia, Nicholas M. Gregg, Gregory A. Worrell, Harvey Huang, Brian N. Lundstrom, Benjamin H. Brinkmann, Tal Pal Attia, Jamie J. Van Gompel, Matt A. Bernstein, Myung-Ho In, John III Huston, Kai J. Miller, Dora Hermes

## Abstract

Stimulation-evoked signals are starting to be used as biomarkers to indicate the state and health of brain networks. The human limbic network, often targeted for brain stimulation therapy, is involved in emotion and memory processing. Previous anatomical, neurophysiological and functional studies suggest distinct subsystems within the limbic network (Rolls, 2015). Previous studies using intracranial electrical stimulation, however, have emphasized the similarities of the evoked waveforms across the limbic network. We test whether these subsystems have distinct stimulation-driven signatures. In seven patients with drug-resistant epilepsy we stimulated the limbic system with single pulse electrical stimulation (SPES). Reliable cortico-cortical evoked potentials (CCEPs) were measured between hippocampus and the posterior cingulate cortex (PCC) and between the amygdala and the anterior cingulate cortex (ACC). However, the CCEP waveform in the PCC after hippocampal stimulation showed a unique and reliable morphology, which we term the limbic H-wave. This limbic H-wave was visually distinct and separately decoded from the amygdala to ACC waveform. Diffusion MRI data show that the measured endpoints in the PCC overlap with the endpoints of the parolfactory cingulum bundle rather than the parahippocampal cingulum, suggesting that the limbic H-wave may travel through fornix, mammillary bodies and the anterior nucleus of the thalamus (ANT). This was further confirmed by stimulating the ANT, which evoked the same limbic H-wave but with a shorter latency. Limbic subsystems have unique stimulation evoked signatures that may be used in the future to help develop stimulation therapies.

**Significance Statement:** The limbic system is often compromised in diverse clinical conditions, such as epilepsy or Alzheimer’s disease, and it is important to characterize its typical circuit responses. Stimulation evoked waveforms have been used in the motor system to diagnose circuit pathology. We translate this framework to limbic subsystems using human intracranial stereo EEG (sEEG) recordings that measure deeper brain areas. Our sEEG recordings describe a stimulation evoked waveform characteristic to the memory and spatial subsystem of the limbic network that we term the limbic H-wave. The limbic H-wave follows anatomical white matter pathways from hippocampus to thalamus to the posterior cingulum and shows promise as a distinct biomarker of signaling in the human brain memory and spatial limbic network.

## Introduction

Describing stimulation evoked biomarkers of specific human brain circuits has greatly advanced the understanding of different brain functions. Studies in the motor system, for instance, described D-waves and I-waves as evoked by direct and indirect excitation (Patton and Amassian, 1954; Awiszus and Feistner, 1994). In other cortical circuits, however, the focus has often been on extracting similar waveforms across connections. Single pulse electrical stimulation (SPES) often evokes negative electrical potential responses in directly connected regions within 50 ms (N1), which has been related to direct cortico-cortical projections (Keller et al., 2014). Recent work has highlighted how the focus on early responses has left out components with different timescales and morphologies (Gronlier et al., 2021; Miller et al.,2021), crucial to unriddling complex cortico-subcortical pathways. Given the important role of the limbic system in neurological diseases, understanding its stimulation-driven features can help advance technologies that target this system.

In 1878, Paul Broca used the term *limbic* (*latin for border*) for the first time to name the brain structures located on the border between cortical and subcortical regions, composed of the cingulate, hippocampal gyri, and the subcallosal frontal area (Bubb et al., 2017). In 1937 connectivity between the hippocampus, mammillary body, anterior thalamic nuclei (ANT), and the cingulate cortex was proposed by James Papez as a functional model for emotions (Papez,1937). Later, in 1949, Paul MacLean built on Papez’s previous work and coined the widely used “Limbic system”, with other cognitive associations (1998). More recent studies have shown that there are multiple subdivisions in the limbic system based on cytoarchitecture with distinct functional roles (amygdala and anterior cingulate cortex (ACC) versus hippocampus and posterior cingulate cortex (PCC)) (Rolls, 2015; Vogt, 2019).

The limbic system also plays a critical role in clinical conditions, such as epilepsy (Bertram et al., 1998; Salanova et al., 2015; Jo et al., 2019), Alzheimer’s disease (Luo et al., 2021), depression (Holtzheimer et al., 2012; Siddiqi et al., 2021) and obsessive-compulsive disorder (Miller et al., 2019; Kahn et al., 2021). In consequence, the limbic network is often targeted with therapeutic brain stimulation to modulate brain function (Lockman and Fisher, 2009; Miller et al.,2019; Gregg et al., 2021). Although therapeutic effects have been shown with hippocampus and ANT stimulation in particular (Lockman and Fisher, 2009; Lozano et al., 2019; Nair et al., 2020; Gregg et al., 2021; Pal Attia et al., 2021), more sophisticated and precise technology has been emerging that senses brain activity in addition to stimulating the brain. Characterizing stimulation evoked waveforms would allow detecting typical or pathological waveforms with such closed-loop systems (Wu et al., 2018).

Characterizing electrophysiological waveforms in the limbic system is relatively challenging with non-invasive (EEG/MEG) techniques due to its deep location. Stereo electroencephalographic (sEEG) electrodes placed during invasive epilepsy monitoring can be used to identify cortical connections of deeper human brain networks (Keller et al., 2014; Enatsu et al., 2015). Direct SPES can be delivered to a particular site while measuring the electrophysiological responses elsewhere (Borchers et al., 2012). Previous studies measuring these cortico-cortical evoked potentials (CCEPs) have confirmed direct anatomical connections within the limbic network using early responses (Matsumoto et al., 2004; Kubota et al., 2013; Enatsu et al., 2015; Oane et al., 2020).

In this study, we show different stimulation-driven waveforms when delivering to and recording from limbic regions (amygdala, HC, ANT and cingulate). We assume that evoked waveforms related to anatomical networks 1) have reliable timing and waveform across trials, 2) share the same features across subjects, 3) white matter endpoints should have reversed polarity across superficial and deeper cortical recording sites, indicating a local current source and sink, and 4) stimulating further downstream in a network should elicit a shorter evoked potential at a recorded endpoint. Using these criteria, we characterize a distinctive limbic H-wave present in HC-ANT-PCC connections, which belong to the hippocampal subsystem of the limbic network.

## Materials and Methods

### Subjects

Data were collected from neurosurgical patients with sEEG probes implanted within the limbic network during invasive epilepsy monitoring. Seven subjects (four males and three females) of ages between 13 and 63 years old (mean 31 years old, Table 1) provided informed consent to participate in the study, which was approved by the Institutional Review Board of Mayo Clinic. During clinical monitoring, the Seizure Onset Zone (SOZ) and regions with interictal activity were identified by epilepsy neurologists [Table 1]. The limbic network was involved in the SOZ only for subject 6, involving a subset of the electrodes in the hippocampus (two out of seven) and posterior cingulate cortex (one out of two) [Table 1, SOZ column].

**Table 1.** Demographic description of subjects included. *SOZ= Seizure Onset Zone; Hc=Hippocampus; PH=Parahippocampal gyrus; Amg=Amygdala; ACC=Anterior Cingulate; MAC=Mid-Anterior Cingulate; MPC=Mid-Posterior Cingulate; PDC=Post-dorsal Cingulate; PVC=Post-Ventral Cingulate; Tha=Thalamus.

| Subject | Age | Sex | Hemisphere implanted | Electrode coverage | SOZ | Inter-ictal notes | Treatment plan |
| --- | --- | --- | --- | --- | --- | --- | --- |
| Sub-01 | 30 | Female | Right | Hc, Amg, ACC, MAC, PDC | Posterior insula/parietal operculum. | Hc | Resective surgery |
| Sub-02 | 19 | Male | Right | Hc, ACC, MPC, Tha | Mid and posterior Insula | Hc | Laser ablation |
| Sub-03 | 31 | Female | Right | Hc, Amg, ACC, MAC, MPC | AC, MAC, and mesial superior frontal gyrus | Hc, inferior mid-frontal orbital gyrus, superior frontal gyrus, white matter | Resective surgery |
| Sub-04 | 13 | Female | Left | Hc, Amg, ACC, MPC, PDC | Inf-posterior insula | Hc, Amg, PDC and medial insula | Laser ablation |
| Sub-05 | 46 | Male | Right | PHc, Amg, MPC, PDC | Sup-frontal gyrus and sulcus, sup-temporal gyrus and sulcus, | PHc, Amg, sup-temporal sulcus, mid-temporal gyrus, mid-temporal lingual gyrus, fusiform gyrus | Implantation of neurostimulator |
| Sub-06 | 63 | Male | Left | PHc, Hc, Amg, PDC, PVC | Anterior and left insula, parieto-temporal region, Hc (2/7) and PDC contacts | Amg, Hc, Inf-parietal supramar gyrus, subcentral gyrus and sulcus. | Diet and medication adjustment |
| Sub-07 | 19 | Male | Left | Hc, Tha, MAC, ACC, MPC, PDC | Medial and Inferior frontal gyrus and sulcus. | Superior, middle, and inferior frontal gyrus, Sup-temporal gyrus and Hc contacts | Resective surgery |

### Electrode localization and inclusion

Multi-contact flexible sEEG probes (DIXI medical) with electrode contacts of 2 mm in length and a diameter of 0.8 mm were implanted. Probes had lengths between 16 mm (5 contacts) to 80.5 mm (18 contacts) [Figure 1B]. The placement of the sEEG probes were selected by the clinical team for the purpose of SOZ localization, with electrode coverage of different brain regions [Figure 1A, sEEG schematic] according to the clinical planning.

**Figure 1.**
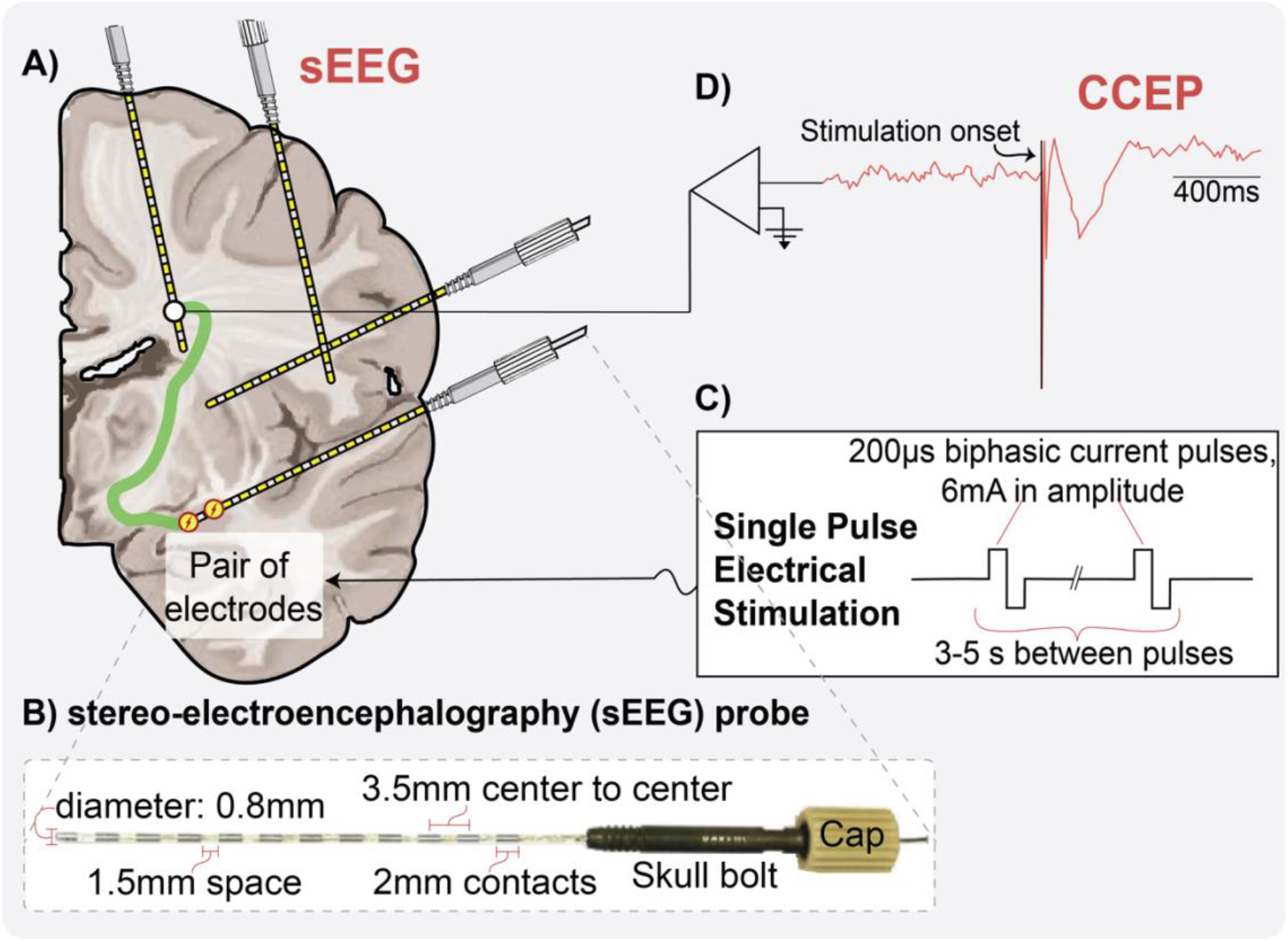
Illustration of Cortico-Cortical Evoked Potentials (CCEPs) experimental paradigm. **A)** Schematic of a brain implanted with sEEG probes. Illustration shows CCEP paradigm where stimulation is delivered and signals depicted in all the other contacts; **B)** Picture of a twelve contacts DIXI medical MICRODEEP sEEG probe with cap and skull bolt and its dimensions; **C)** Parameters used when delivering single pulse biphasic electrical stimulation; **D)** Illustrative schematic of the amplified CCEP signal, showing the stimulation onset followed by a classical evoked potential.

Electrodes were localized using a CT scan (Hermes et al., 2010) and aligned to the T1-weighted (T1w) anatomical MRI using existing software (Friston et al., 2007) (https://www.fil.ion.ucl.ac.uk/spm/software/download/). MRI scans were auto-segmented using FreeSurfer 7 (Fischl, 2012) (https://surfer.nmr.mgh.harvard.edu/). The segmentation was reviewed for accuracy and sEEG electrodes were labeled according to Freesurfer’s Destrieux atlas (Destrieux et al., 2010). Electrodes labeled as amygdala, hippocampus, parahippocampal gyrus, thalamus, and cingulate cortex were included in the study. Sites labeled as *anterior cingulate* and *middle anterior cingulate* were grouped together as Anterior Cingulate Cortex (ACC); and sites labeled as *middle posterior cingulate, posterior-dorsal cingulate*, and *posterior-ventral cingulate* were grouped together as Posterior Cingulate Cortex (PCC). Sites labeled as *hippocampus* and *parahippocampal gyrus* were grouped together as *Hippocampal Complex (HC) (Nemanic et al., 2004)*. Estimated positions of the electrodes are shown in Supplemental Figures 2-1 and 2-2.

### CCEPs and Intracranial EEG measurements

Using a Nicolet Cortical Stimulator from Natus Medical Incorporated (Middleton, WI). SPES was delivered using biphasic pulses of 200μs duration. For subjects 1 to 6, SPES was applied with 6mA amplitude every three to five seconds for a total number of 10 to 12 times [Figure 1C]. Given the higher excitability observed in subject 7, lower amplitudes (3mA and 4mA) were used for most sites. CCEPs were recorded [Figure 1D] at 2048Hz using a Natus Quantum amplifier.

### Data pre-processing

CCEP recordings were visually inspected for electrical and movement artifacts, and channels and trials with excessive line noise were excluded from analyses. Data were re-referenced to a modified common average in a trial-by-trial manner to exclude the stimulated channels and channels with large variance. In addition, we consider the fact that noise was shared across blocks of 64 channels because 64 channels were acquired within one headbox (maximum of 4 headboxes with 256 channels total). For each channel, we consider the 64 channels from the same headbox to calculate a common reference signal. From the 64 channels, we exclude bad channels, stimulated channels, the 5% of channels with the largest variance from 500-2000 ms after stimulation and the 25% of channels with the largest variance from 10-100 ms after stimulation to calculate the average. The common average was then subtracted from all other channels. A baseline correction was then used by subtracting the median amplitude from 0.5 s to 0.05 s before each stimulation from each trial.

### Statistical analyses

#### Reliable CCEP waveforms and significance

One of the criteria we assume for waveforms related to anatomical networks is that they have a reliable waveform across trials. To determine whether responses were significant, we therefore tested the reliability of the shape of the responses across trials using a previously published method (Miller et al., 2021). Electrodes were stimulated 10-12 times, and a cross projection matrix, P, was calculated for each stimulation electrode pair as scalar projections between all pairs of trials from 15-500 ms, with self-projections removed: 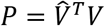, where 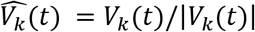 for the k^th^ trial. Significance can then be assessed by calculating a t-statistic and associated p-value testing whether the cross projections in the matrix *P* differ significantly from zero. The advantage is that this method fully depends on inter-trial reliability, while not depending on the shape of the CCEP and no prior assumptions are made for response duration or polarity. For multiple comparison correction across all stimulation and response pairs, we used FDR-adjusted *p*-value <0.05.

#### Time-lag between inputs from different sites

In the limbic circuit, the PCC is connected to the HC and ANT. To understand whether PCC responses to HC and ANT stimulation were similar, but lagged in time, we calculated time-lagged cross-correlations between HC and ANT evoked responses in PCC within two subjects that had electrodes placed in all three areas. Cross correlations were calculated between all trials with PCC responses when the HC was stimulated (***n*** trials) and when ANT was stimulated (***m*** trials), resulting in ***m*** x***n*** cross correlations. To calculate confidence intervals of the cross-correlation and time-lag between the two inputs, we used a bootstrapping method (Efron and Tibshirani, 1998) and sampled with replacement 10,000 times from observed cross-correlations.

#### Wavelet Principal Component Analysis (PCA) and Linear Discriminant Analysis

One of the criteria we assume for waveforms related to anatomical networks is that they have a reliable waveform across subjects. We quantitatively assess distinct waveforms in the amygdala and hippocampal limbic subsystems across subjects. PCA was performed in the discrete wavelet domain on significant CCEPs between limbic regions to provide a low-dimensional visualization of (dis)similarity between tested limbic connections. Subject 1, with both amygdala-to-ACC (n=9) and HC-to-PCC CCEPs (n=15), was withheld as independent testing data. First, the significant CCEPs between limbic regions were L2-normalized between 100 and 1000 ms post-stimulation. Then, the highest level (8) discrete wavelet transform of each CCEP was calculated using the fourth Symlet wavelet, and all coefficients below the 95th percentile were set to 0. The wavelet transformation and thresholding steps allow rapid temporal changes in the signal to be emphasized while reducing low-amplitude noise (Daubechies, 1992; Gupta and Jacobson,2006; Puyati et al., 2006; Brunton and Kutz, 2022). The set of all transformed CCEPs was mean-centered at each wavelet coefficient and principal components were determined by singular value decomposition. CCEPs were projected to two principal components that independently showed good separation between the amygdala-to-ACC and HC-to-PCC conditions, by visual inspection. Linear discriminant analysis was performed in this two-dimensional space between those two limbic conditions using leave-one-subject-out training, and model accuracy was later assessed.

### Diffusion-weighted imaging (DWI) and tractography

Subjects 2 and 7 were scanned in a Compact 3.0T MRI scanner with high-performance gradients (Foo et al., 2018) at Mayo Clinic Rochester under an IRB-approved protocol. We used Distortion-free imaging: A double encoding method (In et al.) to scan two series with each two volumes at b=0 s/mm^2^ and 48 directions at b=1000 s/mm^2^, TR/TE/TE_NE_ = 2659/42.7/49.6 ms (TE_NE_ is navigator echo time for the DIADEM sequence), 70 slices at 2 mm thickness (zero gap), FOV of 216 mm and acquisition matrix 108X108.

Diffusion Magnetic Resonance Imaging (dMRI) data were preprocessed to correct for subject motion and eddy currents and to align the dMRI images and T1w anatomical image using the ANTs (Advanced Normalization Tools) algorithm in QSIprep version 0.14.2 (Cieslak et al., 2021). DSI studio was used to track different subcomponents of the cingulum bundles and the fornix bundles.

First, the restricted diffusion was quantified using restricted diffusion imaging (Yeh et al., 2017). Next, the diffusion data were reconstructed using generalized q-sampling imaging (Yeh et al.,2010) with a diffusion sampling length ratio of 1.25. Finally, a deterministic fiber tracking algorithm (Yeh et al., 2013) was used with augmented tracking strategies (Yeh, 2020) to improve reproducibility. The anatomy prior of a tractography atlas (Yeh et al., 2018) was used to map the fornix and cingulum bundles with a distance tolerance of 16 (mm). The anisotropy threshold was randomly selected, and the angular threshold was randomly selected from 15 degrees to 90 degrees, the step size was randomly selected from 0.5 voxel to 1.5 voxels. Tracks with lengths shorter than 20 mm or longer than 300 mm were discarded. A total of 5000 tracts were calculated. Topology-informed pruning (Yeh et al., 2019) was applied to the tractography with 12 iteration(s) to remove false connections.

### Code and data accessibility

Code written in MATLAB to reproduce the statistics and figures contained in this manuscript will be available on our GitHub page. Data will be shared with publication in BIDS format on OpenNeuro.org.

## Results

In order to understand whether limbic subsystems have distinct stimulation evoked network signatures, we stimulate and measure from different limbic regions. We first visualize the waveforms across subjects [Figure 2], showing various waveforms across connections and highlighting the distinct connectivity between ACC and the amygdala versus PCC and the hippocampal complex (HC). Second, we show how reliable the HC to the PCC (HC-to-PC) waveform is across trials in one subject, and that it displays a characteristic waveform, which we will call the *limbic H-wave* [Figure 3]. Third, we show the HC-to-PCC waveforms across all subjects and connections, typically showing the *limbic H-wave* with a characteristic peak at 200 ms. The proportion of reliable waveforms from the HC-to-PCC is higher than from the HC-to-ACC or amygdala-to-PCC, further establishing the strength of connectivity between HC and PCC [Figure 4]. Fourth, to confirm the current source and sink of the *limbic H-wave* peak at 200 ms in the PCC, we show that the peak is typically reversed at deeper and superficial PCC recording sites [Figure 5]. Fifth, we show that stimulating more posterior in the HC evokes faster responses in the PCC [Figure 6]. Sixth, since a 200 ms delay is too long for a direct connection, we further probe the network involved in these PCC waveforms by stimulating the ANT and extracting the white matter bundles potentially involved in the propagation of the waveform in 2 subjects [Figure 7]. Lastly, to understand the decodability between amygdala and ACC versus hippocampus and PCC across subjects, we test whether the waveform shape could be predicted from existing subjects in a left-out-subject [Figure 8].

**Figure 2.**
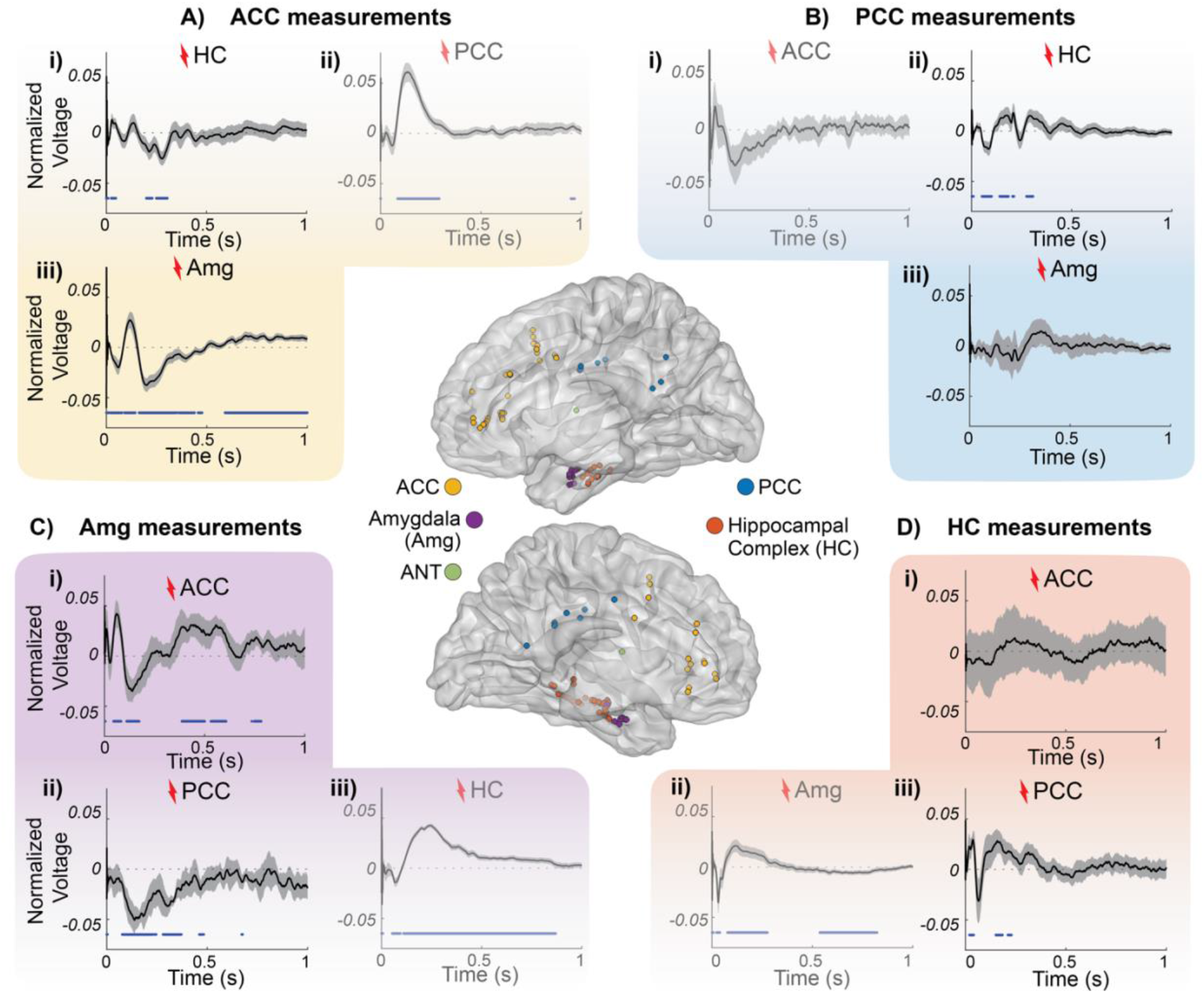
Stimulation-driven waveforms in limbic connections across subjects. MNI brain render of both right (upper) and left (bottom) hemispheres with all limbic contacts where signals from A, B, C, and D were measuring and stimulating from. Normalized voltage of significant CCEPs (thicker black line) plotted over time with confidence interval (gray shadow) and samples with a significant difference from baseline (dots at the bottom of plots). **A)** Measurements in the ACC after HC (i), PCC (ii), and amygdala (iii) stimulation. **B)** Measurements in the PCC after ACC (i), HC (ii), and amygdala (iii) stimulation. **C)** Measurements in the amygdala after ACC (i), PCC (ii), and HC (iii) stimulation. **D)** Measurements in the HC after ACC (i), amygdala (ii), and PCC (iii) stimulation.

**Figure 3.**
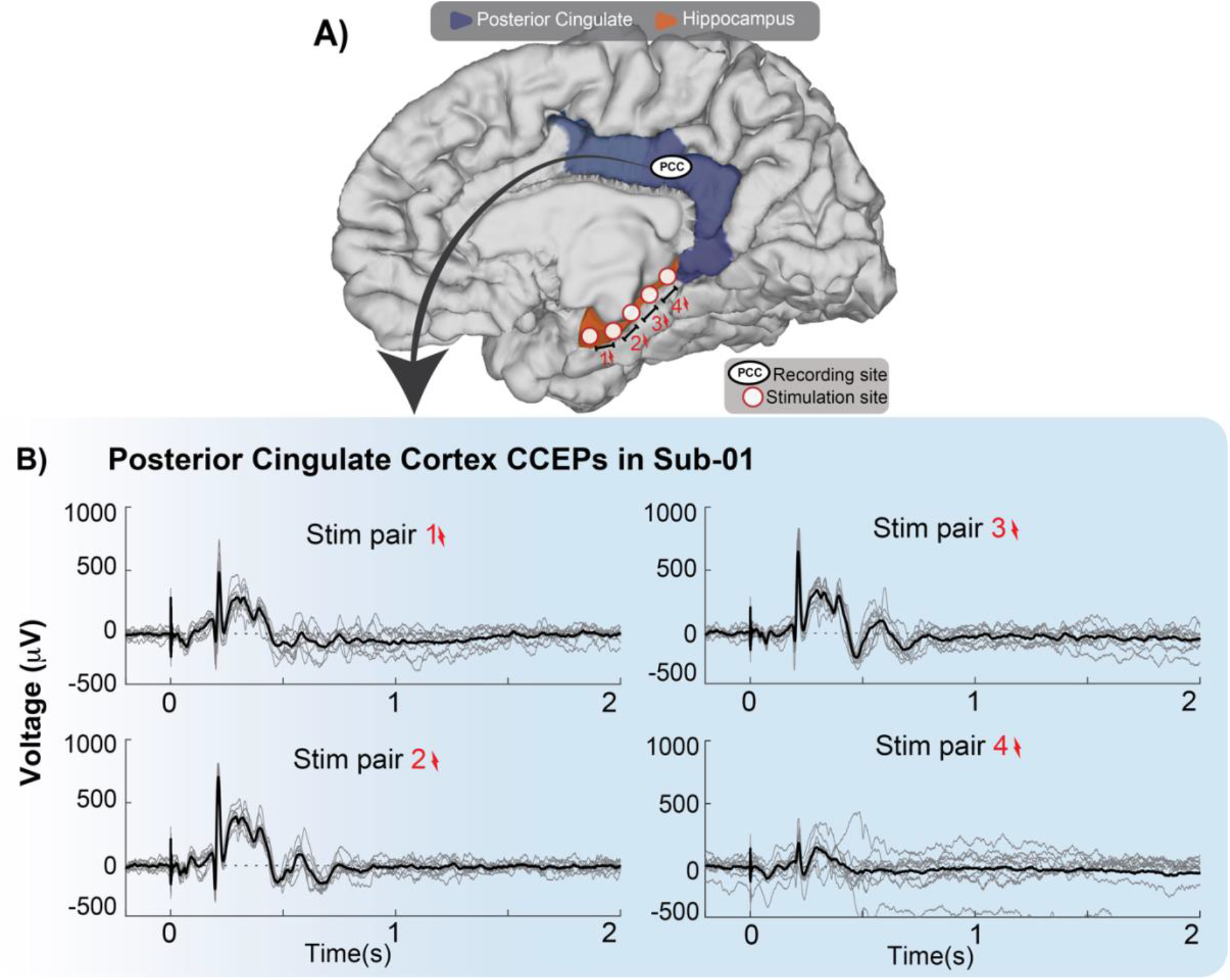
PCC shows a reliable late and complex waveform when stimulated along the hippocampus (limbic H-wave). **A)** Brain schematic with representative recording and stimulation sites, with PCC in blue and areas surrounding hippocampus in orange since hippocampus is located underneath the cortical surface. **B)** CCEPs (μV) from a single contact in PCC from subject one. Each panel shows CCEP trials (gray lines) after stimulating a different pair of hippocampal electrodes. Stimulation onset at time 0 and the average response is shown in black.

**Figure 4.**
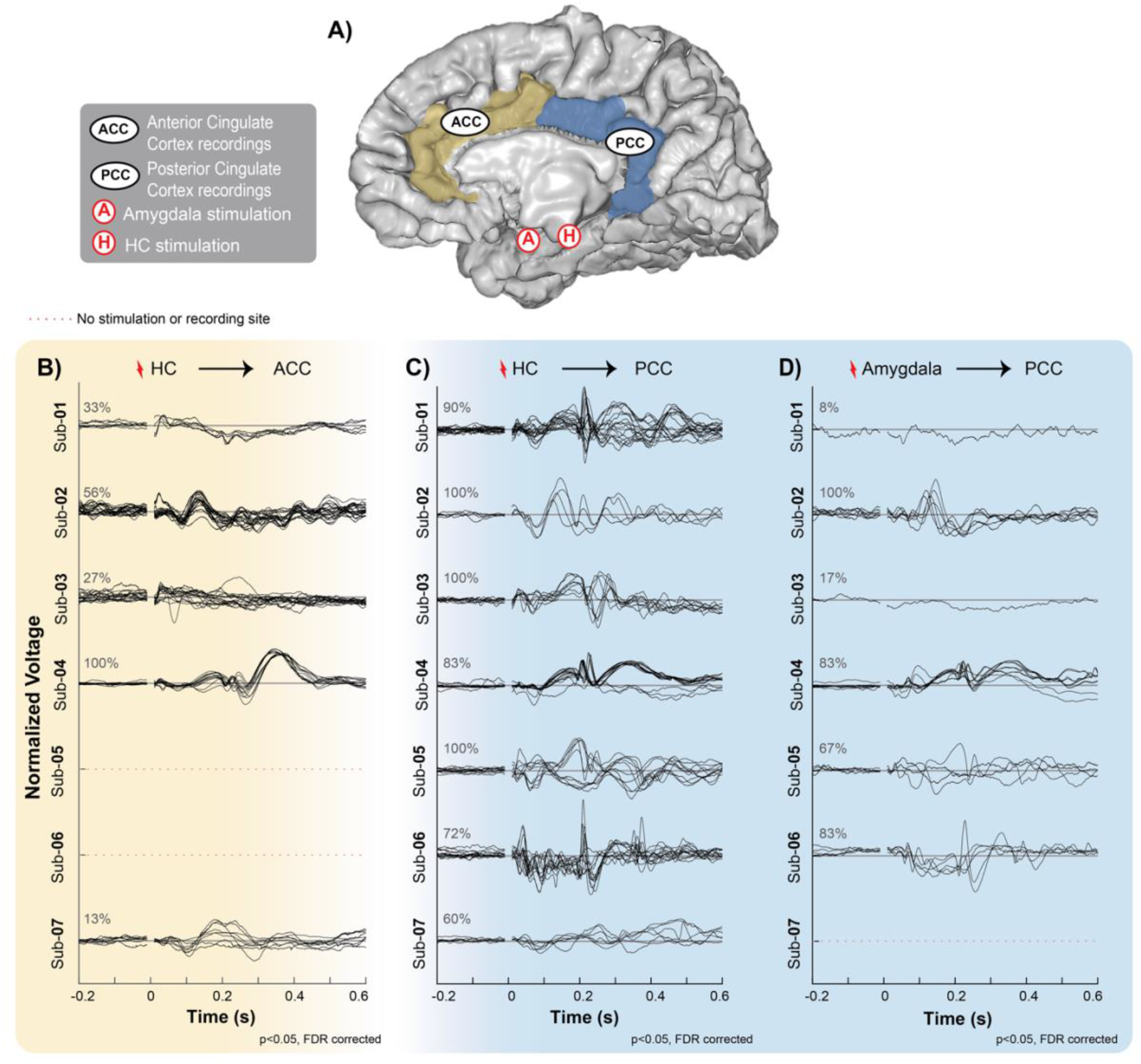
CCEPs measured from PCC and ACC after hippocampal and amygdala stimulation. **A)** Brain schematic with representative recording and stimulation sites. Gray matter labeled as PCC (blue) and ACC (beige) contain an example of a recording site. (A) = Amygdala and (H) = Hippocampal complex sites representing the location of the stimulating electrodes. Significant average responses (t-tests, p_FDRcorrected_<0.05, L2-normalized) are represented by each black line under three different conditions, denoted in B), C) and D). **B)** ACC under hippocampal stimulation, **C)** PCC under hippocampal stimulation, and **D)** PCC under amygdala stimulation.

**Figure 5.**
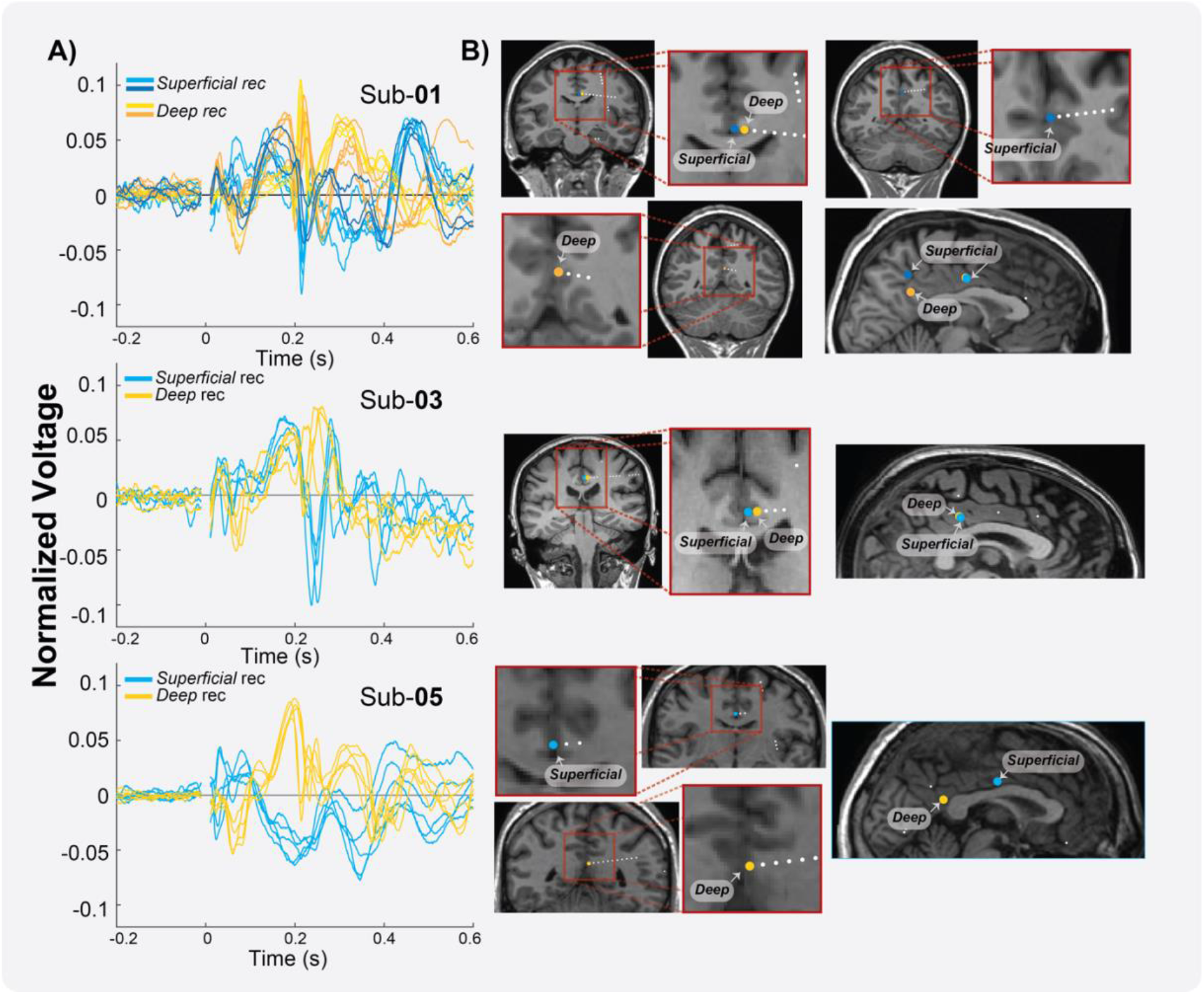
Electrodes positioned in different depths of the PCC and variability in morphology. **A)** Normalized limbic H-waves (L2-normalized) in the PCC from hippocampal stimulation plotted as a function of time. Limbic H-waves from a superficial electrode (blue traces), limbic H-waves from the deep electrode (yellow traces). **B)** Coronal view (left panel) and sagittal view (right panel) showing an estimation of the electrodes (blue, superficial; yellow, deep) for subject one, three, and five.

**Figure 6.**
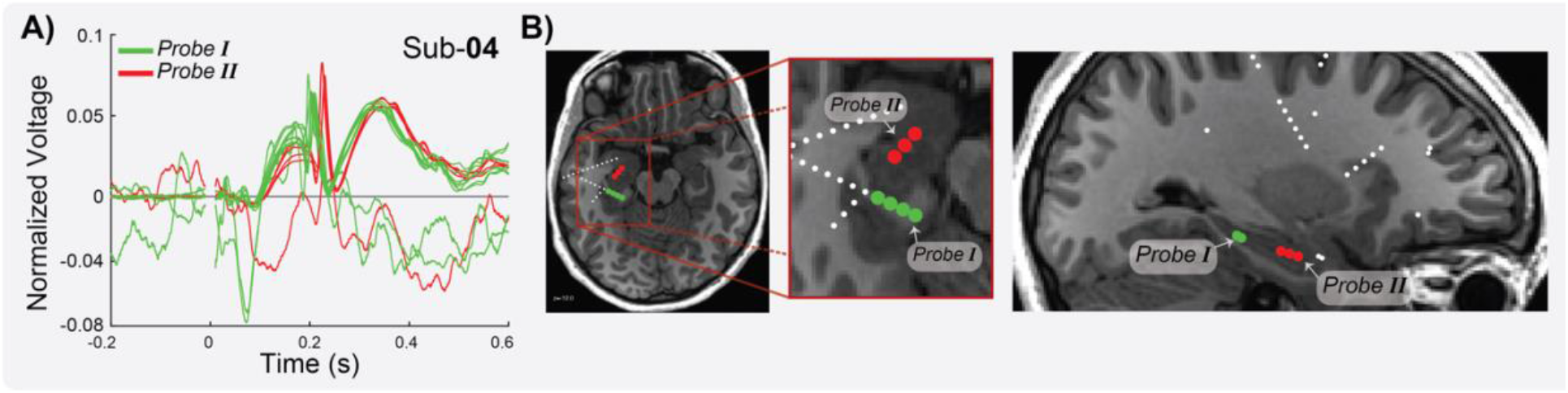
Location of stimulation sites and PCC response. **A)** Normalized limbic H-waves (L2-normalized) measured in PCC after hippocampal stimulation are plotted as function of time for subject 4. Significant limbic H-waves upon stimulation of Probe I (in green) and Probe II (in red). **B)** Axial (left) and sagittal view (right) with an estimation of the stimulating electrodes (contacts in Probe I in green; contacts in Probe II in red).

**Figure 7.**
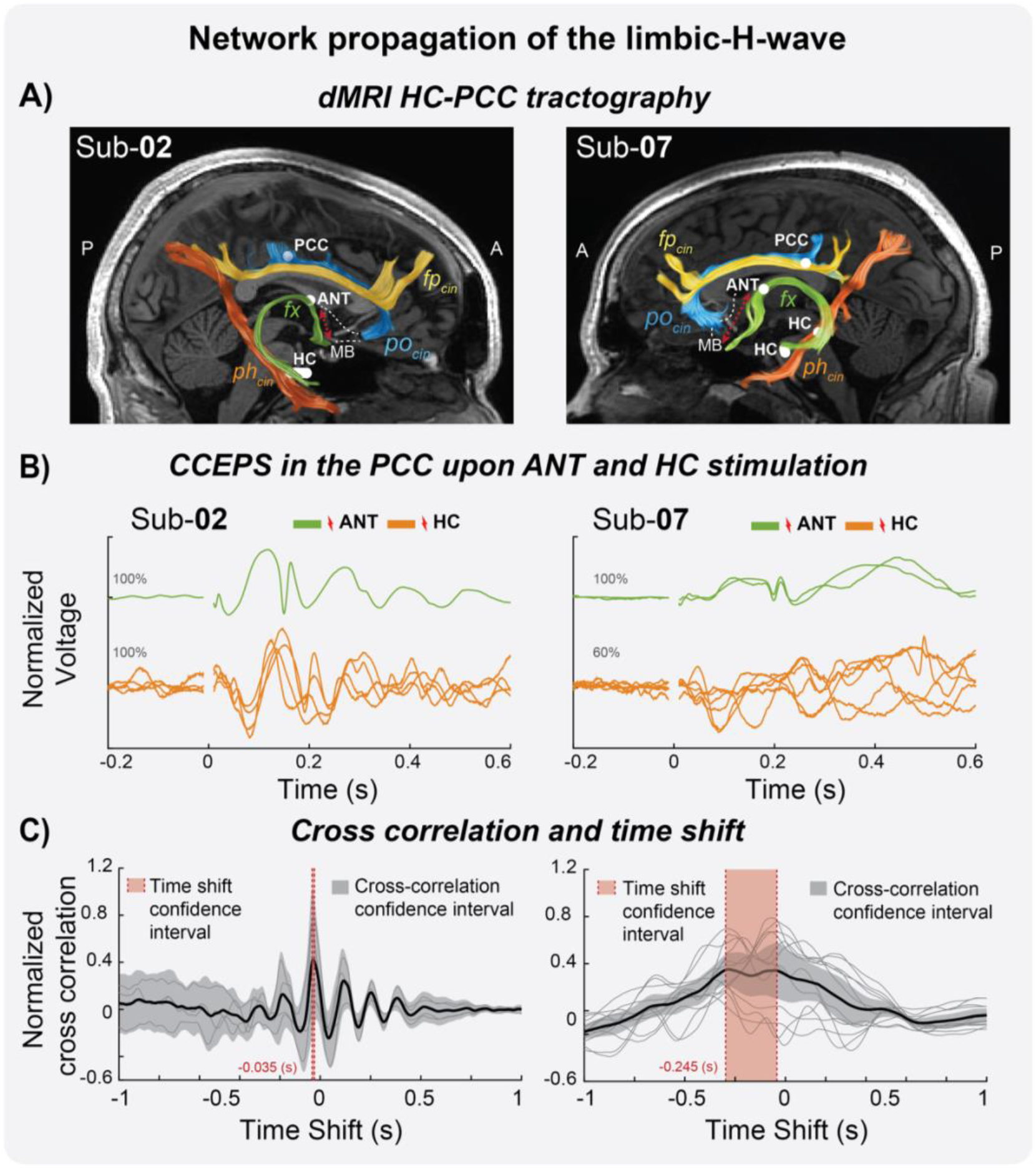
Limbic H-waves under ANT stimulation with reduced latencies reveal an anterior propagation route. **A)** dMRI estimates of the fornix (**fx**, red tract), parahippocampal (**ph_cin_**, orange tract), frontoparietal (**fp_cin_**, yellow), parolfactory (**po_cin_**, blue) cingulum, and estimated location of the recording and stimulation sites in the PCC, HC and ANT (white circles). Additionally, the representative MTT (dotted red line) connecting the mammillary body (MB) and the ANT. **B)** L2-normalized PCC CCEPs under ANT (green) and Hippocampal (orange) stimulation, plotted over time. **C)** Normalized cross-correlations over time-shift in seconds. Single cross-correlations (gray) and their average (black, 95% confidence interval, transparent gray). CCEPs in the PCC time-shift after ANT stimulation compared to HC stimulation (95% confidence interval, transparent red).

**Figure 8.**
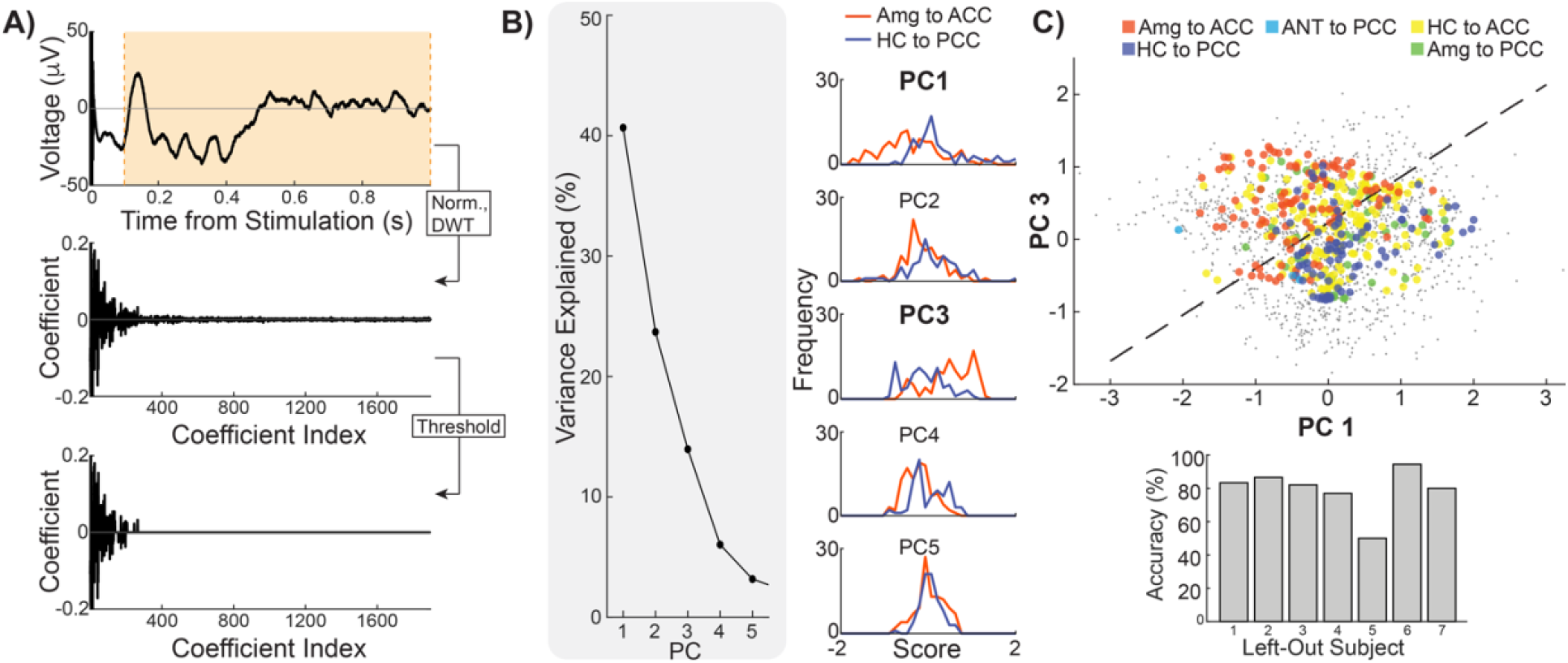
Similarity metric within tested limbic connections. **A)** Each significant limbic CCEP is L2-normalized between 100 and 1000 ms post-stimulation and then subjected to a level-8 (maximum) discrete wavelet transform using the fourth Symlet wavelet. All wavelet coefficients below the 95% percentile were set to 0. **B)** Principal component analysis was performed on all thresholded and discrete wavelet-transformed CCEPs. (Left) percent variance explained by each of the first five principal components. (Right) Distribution of amygdala-to-ACC and hippocampus-to-PCC scores in each of the first five principal components. **C)** (Top) All limbic CCEPs were projected to the first and third principal components. Each color representing a different indirect limbic connection, and gray dots representing waveforms of other limbic connections (e.g., within regions, such as HC-to-HC [supplemental figure 2-3] or amg-to-amg [supplemental figure 2-4], and connections of adjacent regions, such as Amg-to-HC [supplemental figure 2-3]) Dashed line indicates decision boundary for linear discriminant analysis between amygdala-to-ACC and HC-to-PCC CCEPs. (Bottom) Leave-one-subject-out cross validation yields a mean accuracy of 79%.

### Limbic network waveforms across subjects

First, we visualize the stimulation-evoked network signatures of the different limbic connections across all subjects. Figure 2 shows the CCEP waveform averaged across all significant responses (p_FDRcorrected_<0.05, see methods) across all subjects [Figure 2, middle panel with standard brain shows the coverage across all subjects. Supplemental Figure 2-1 and 2-2 show the estimated coverage for each subject]. A broad range of stimulation-driven waveforms is observed throughout the limbic network. When stimulating and measuring most adjacent sites (PCC to ACC [Figure 2A], HC to Amygdala [Figure 2C], Amygdala to HC [Figure 2D]), evoked potentials significantly differ from zero across earlier and later periods, showing early waveforms with a peak followed by a slow wave [Figure 2-3 and 2-4 show measurements in HC and amygdala in each individual subject]. Between distant sites, measurements in the ACC show significant waveforms from amygdala stimulation of large amplitude, starting early and lasting up to 1 sec duration, whereas hippocampal stimulation caused smaller deviations [Figure 2A]. Measurements in the PCC show significant responses from HC stimulation with an early peak before 100 ms and a later peak around 200 ms [Figure 2B]. This 200 ms peak is a much later response than the short N1 latencies typically reported within the first 100 ms. Measurements in amygdala show significant changes across the subjects from ACC or PCC stimulation. Measurements in hippocampus however show significant evoked waveforms up to 500 ms only from PCC stimulation. While these data emphasize the strong bidirectional connections between hippocampus and PCC or amygdala and ACC, the averaging across subjects may obscure some of the details in the waveforms in terms of reliability across trials, temporal shifts and polarity.

### CCEPs in PCC after HC stimulation

To ensure these unique features in the HC-to-PCC waveform are also present across trials, we show data from subject 1 [Figure 3]. After stimulating different pairs of electrodes located along the HC, we analyze the data from a single PCC site [schematic in Figure 3A]. In this example, prominent and very consistent CCEPs are observed in the PCC when stimulating along HC electrode-pairs [Figure 3B]. Responses in all trials show a sharp peak around 200 ms, that is preceded and followed by slow positive waves until about 500 ms. Thus, we confirm that the waveform is reliable across trials and is not driven by outliers or fluctuating interictal activity. We will further refer to this waveform as the *limbic H-wave*.

To further observe the presence of the limbic H-wave in different electrodes on HC to PCC connections, we show the CCEP waveforms between all significant stimulated/measured connections [Figure 4C]. Significance was established by cross-trial reliability and we find that from all possible HC-to-PCC connections across subjects, a large percentage of connections are significant (87%, range 60-100%, t-tests, p_FDRcorrected_<0.05). We find that from all possible HC-to-ACC connections across subjects, fewer connections are significant (46%, range 17 - 100%, t-tests, p_FDRcorrected_<0.05) [Figure 4B]. Amygdala-to-PCC connections are also less reliably observed across subjects (60%, range 8-100%, *t*-tests, p_FDRcorrected_<0.05) [Figure 4D] (voltage IS normalized in Figure 4, Supplemental Figure 4-1 shows the CCEPs in μV). These calculations confirm the strong connectivity between HC and PCC in each individual subject.

While all shown waveforms are reliable across trials, these reliable waveforms show some variability between connections (Figure 4B). A common feature observed in most HC-to-PCC connections is a sharp peak around 200 ms, except for subject 7 (we note that stimulation amplitude was reduced in subject 7). Evoked waveforms are often positive in polarity with a total duration of around 500 ms. Differences in polarity and latency will be addressed in subsequent Figures 5 and 6. Importantly, these responses show more complex features than previously described in studies focusing on the early N1/N2 components.

### Polarity of limbic H-wave reverses at PCC endpoints

Polarity reversals between contacts recording from a superficial and a deep source across the cortical surface indicate that there is a signals source between the contacts (Mitzdorf, 1985). We observe that HC-to-PCC connections often elicited a limbic H-wave with a sharp peak around 200 ms and an overall latency of around 500 ms. However, the peak could be either positive or negative (Figure 4C). We hypothesize that the variability in polarity is a result of sEEG electrodes recording from different cortical depths. Therefore, we inspect the signals from subjects 1, 3, and 5 where both polarities are present. Figure 5A shows limbic H-wave from different measurement electrodes in the PCC. Differences in polarity are observed in *s*ignals recorded from superficial sites [Figure 5, dark and light blue traces in *A;* blue circles representing electrodes in *B*] compared to signals depicted deeper in the gray matter (around 3.5 mm center to center) [Figure 5, dark and bright yellow traces in *A*; yellow circles representing electrodes in *B*]. The superficial recordings show a peak with negative polarity around 200 ms, whereas the deeper recordings show a peak with positive polarity. These visual inspections show that the position of the electrodes relative to the gray matter determines the polarity of the limbic H-wave, indicating a local signal source and sink in the PCC.

### Variability in latency of responses

To further understand whether the waveform is related to the HC-PCC subsystem of the limbic network, we test whether stimulation at posterior sites along the HC elicits earlier evoked responses in the PCC. We observe that hippocampus to PCC connections often elicited a limbic H-wave with a sharp peak around 200 ms, but the timing could differ between stimulation and recording pairs. We hypothesize that the variability in timing is a result of sEEG electrodes positioned at different distances, as some signals travel further than others. In subject 4, we group together limbic H-waves elicited by single sEEG probes within the HC, ending up with a posterior *(Probe I)* [Figure 6B, *green circles]* and an anterior group *(Probe II)* [Figure 6B, *red circles*], and visualize the limbic H-waves in different colors (green for posterior stimulation, red for anterior stimulation) [Figure 6A]. Stimulation in more posterior HC sites elicits earlier peaks (green traces) than stimulation of more anterior HC sites (red traces).

We note that responses from stimulation of more posterior HC sites are about 25 ms earlier compared to sites that are 1 or 2 cm more anterior. This difference in latency is unlikely to be related only to the distance along the fornix, given that typical white matter transmission speeds are much faster than 40 cm/sec (Innocenti et al., 2014). However, the longitudinal axis of the hippocampus contains different sub-fields (Strange et al., 2014) which may play a role as well. This variability in latencies suggests that signals from the HC-to-PCC spread in an anterior-to-posterior direction along the HC.

### ANT stimulation elicits similar PCC waveforms, with reduced latencies

Diffusion MRI (dMRI) imaging has delineated several subsegments of the cingulum bundle. There are two possible pathways from the HC to the PCC, the posterior and the anterior. The posterior, connecting through the parahippocampal cingulum bundle (ph_cin_) [Figure 7A, orange tract] (Jones et al., 2013; Wu et al., 2016); and the anterior, connecting through the thalamic portion of the limbic system, where hippocampus projects through the fornix (fx), to the mammillary bodies (MB), which project through the mammillothalamic tract (Grewal et al., 2018) to the ANT, that further projects to the PCC through the parolfactory cingulum (po_cin_) bundle (Mufson and Pandya, 1984; Jones et al., 2013; Wu et al., 2016; Wang et al., 2020; Gregg et al.,2021; Aggleton et al., 2022). DSI studio software tool can separately estimate different subsegments of the cingulum bundle (Wu et al., 2016). Figure 7A shows two subjects with the estimates of the fornix, as well as different cingulum subsegments: the frontoparietal (fp_cin_) [yellow tract], ph_cin_[orange tract], and po_cin_,[blue tract], as shown in previous anatomical studies (Jones et al., 2013; Wu et al., 2016; Bubb et al., 2018). In addition, estimates of the electrode sites [Figure 7, white circles] in the HC, the ANT, and the PCC.

We analyzed CCEP data from subjects 2 and 7 with electrodes implanted in the ANT trying to differentiate the pathway potentially involved in the propagation of the limbic H-wave. From all possible ANT-to-PCC connections, a large proportion are significant (84%, range 67-100%), where the presence of limbic H-waves is observed with the characteristic 200 ms peak [Figure 7B, green traces - Figure S2C]. To quantify the delay between the limbic H-waves measured in the PCC after ANT and HC stimulation, we calculated the cross-correlation between the average of the HC-to-PCC and ANT-to-PCC limbic H-waves [Figure 7C]. The ANT-to-PCC connections are significantly faster compared to the HC-to-PCC limbic H-waves (confidence intervals calculated by bootstrapping with 10,000 resamples), with an average shift of 34 ms in subject 2 and 245 ms in subject 7. The 34 ms observed in subject 2 is a typical delay between distant regions (Keller et al., 2014), the 245 ms in subject 7 also has a larger confidence interval give that the hippocampal stimulations at 4mA are less robust. The decreased latency of the limbic H-wave after ANT stimulation suggests an anterior route of propagation, potentially traveling from HC, through the fornix to the ANT, and later through the parolfactory cingulum bundle to the PCC, (Wu et al., 2016; Wang et al., 2020).

By looking at the visual match between the electrode positions, estimated bundles and their endpoints, in addition to the presence of the limbic H-wave after ANT stimulation, we hypothesize that the limbic H-wave is generated by the circuit between the hippocampus, ANT, and the PCC, consistent with the previous reports on the hippocampal limbic system (Rolls,2015; Wang et al., 2020; Aggleton et al., 2022). Interestingly, limbic H-waves in the PCC are not as clearly observed in subject 7 under HC stimulation (Figure 7A, orange traces), whereas ANT stimulation evoked clear limbic H-waves. We note that the stimulation amplitude in subject 7 was lower compared to the other subjects (4mA compared to 6mA). Thus, a lower amplitude results in a smaller volume of tissue activation, perhaps not stimulating the hippocampus or fornix as strongly as in other cases.

### Limbic connections

We report the limbic H-wave, an electrophysiological signature in the PCC when stimulating in the HC. This limbic H-wave is visually distinct from other waveforms, including those with strong effective and functional connectivity, such as the amygdala-to-ACC (Beckmann et al., 2009;Rolls, 2015; Bubb et al., 2018; Oane et al., 2020). To quantify the distinction between these two functional networks: memory and spatial HC-to-PCC and the emotional amygdala-to-ACC connections, we performed a linear discriminant analysis on principal components of discrete wavelet-transformed CCEPs (see methods) [Figure 8]. The first and third principal components, which collectively explained 55% of total variance across CCEPs, were used for linear discriminant analysis as they independently showed good separation between HC-to-PCC and amygdala-to-ACC conditions [Figure 8B, right panel]. Figure 8C (top) shows that HC-to-PCC CCEPs (blue) cluster distinctly from amygdala-to-ACC (red) CCEPs. The mean leave-one-subject-out cross validation accuracy of linear discriminant analysis between these two conditions is 79% (Figure 8C, bottom), much higher than the chance level of 50%. The other limbic conditions are more interspersed in this two-dimensional representation.

## Discussion

In order to map network signatures in limbic subsystems, we measured single pulse stimulation-evoked electrophysiological waveforms from the human limbic system across the full duration of the response (1 sec), well beyond previously characterized early responses (<100 ms). Our data showed different stimulation-driven waveforms when stimulating and recording from different limbic regions. We describe how the limbic H-wave, measured in PCC after hippocampal stimulation: 1) shows reliable timing and morphology across trials with a peak around 200 ms, 2) shares the same, decodable, features across subjects, 3) has reversed polarity across superficial and deeper cortical PCC recording sites, and 4) has a decrease in latency of the response at a recorded endpoint when stimulating further downstream in the hippocampus or in the ANT. Following these criteria, we characterize a distinctive limbic H-wave present in indirect HC-ANT-PCC connections that is likely related to the memory and spatial hippocampal subsystem of the limbic network.

### Responses in the memory and spatial limbic subsystem

Using the limbic system as a canonical circuit for investigation demonstrates that CCEPs can map out network signatures in cognitive brain circuits, such as the hippocampal limbic subsystem associated with memory and spatial processing. Our findings indicate that HC-to-PCC connections have a distinct waveform with a sharp peak around 200 ms preceded and followed by slow waves. These late responses are likely generated through polysynaptic indirect cortico-subcortical connections (Child and Benarroch, 2013; Kubota et al., 2013; Kumaravelu et al., 2018). Brain structures involved in the propagation of the limbic H-wave, such as hippocampus, ANT and PCC, have been described as part of the Papez circuit (Papez, 1937), and later associated with a distinct functional role in memory and spatial processing within the limbic system (Rolls, 2015; Bubb et al., 2017).

The dMRI data indicate the white matter tracts that potentially mediate the propagation of the limbic H-wave. The stimulated and recorded electrode sites matched the fornix and parolfactory cingulum bundle endpoints respectively. Diffusion MRI studies have started to delineate different parts of the cingulum bundle in humans (Beckmann et al., 2009; Jones et al., 2013; Wu et al., 2016). The parolfactory cingulum bundle connects subgenual regions to the PCC, consistent with what has been shown in animal studies (Mufson and Pandya, 1984; Carmichael and Price, 1996; Wu et al., 2016; Bubb et al., 2018). Given the endpoint location of our electrodes, the po_cin_ segment of the cingulum is most likely involved in the limbic H-wave, in contrast with the fp_cin_ and ph_cin_ segments of the cingulum, where the endpoints are located more posterior from the electrodes. Further animal, lesion, and stimulation studies may further explain the network involved in the limbic H-wave and how its later components are being generated.

It is important to distinguish the connections from amygdala and ACC versus the hippocampus and PCC to understand limbic subsystems (Rolls, 2015). Recent functional studies further emphasize the subdivisions of the cingulate cortex and cytoarchitectural differences (Aponik-Gremillion et al., 2022; Willbrand et al., 2022). We trained a model that showed a distinction between HC-to-PCC (related to the hippocampal limbic system) and amygdala-to-ACC connections (related to the emotion limbic system) based on its waveform. This indicates that electrical stimulation evoked waveforms are particular for different anatomical connections and may potentially serve as biomarkers of different anatomical and functional limbic subsystems, in a similar manner as other studies have done in the motor system.

The limbic H-wave in our data is consistently observed across subjects. However, variability within subjects shows how unique electrophysiological responses vary in amplitude, polarity, and latency depending on the stimulated and measured electrode. The high temporal and spatial resolution of the sEEG recordings described here allow us to delineate how shifts in polarity and latency can be explained by the cortical layer of the recording electrode and the distance between the recording and stimulated site.

Previous CCEP studies have primarily described the early evoked responses by focusing on 1) changes in strength (Root Mean Square, RMS), or on 2) the first negative (N1) or positive (P1) components (Kubota et al., 2013; Enatsu et al., 2015; Donos et al., 2016; Takeyama et al.,2019; Oane et al., 2020). Kubota and Enatsu reported strong CCEPs in the PCC after both anterior and posterior hippocampal stimulation. While their results thus focus on the early responses, a review of their figures reveals a similar waveform within the first 300 ms that is similar to the limbic H-wave in our data. Another study similarly showed a peak in the PCC at 187 ms after fornix stimulation (Koubeissi et al., 2013). While neither study elaborated upon this waveform, they provide independent confirmation of our measurements. Thus, the limbic H-wave in our data is therefore reproducible across studies and emphasizes that different networks may show unique interactions.

### Stimulation driven signatures as biomarkers

In motor systems, Patton and Amassian described D-waves as early or direct responses, in contrast with the later or indirect I-waves (Patton and Amassian, 1954). These well characterized waveforms are used as biomarkers for intraoperative monitoring, to understand and diagnose pathology in the motor functions (Boyd et al., 1986; Hicks et al., 1991; Quinones-Hinojosa et al., 2005; Costa et al., 2013). In the limbic system, such waveforms thus far have not been characterized. The limbic H-wave we observed may characterize an indirect anterior route within the limbic network (Papez, 1937; Rolls, 2015), from hippocampus, through the fornix to the mamillary bodies, through the mammillothalamic tract to ANT, through the parolfactory segment of the cingulum bundle to the PCC [Figure 7].

Epilepsy often involves the limbic system (Wyllie, 2012). In our study, only subject 6 had a limbic SOZ (see methods, subjects), and only subject 4 had reported interictal activity in the PCC [Table 1, inter-ictal notes]. However, all 7 subjects had reported interictal epileptiform activity involving the hippocampus or amygdala, which is typical for patients who have electrodes implanted in the limbic system. Interictal epileptiform spike-slow wave discharges have been reported in the cerebral cortex in animals behaving freely (Pearce et al., 2014), and is typically considered a hallmark of hyperexcitability associated with epilepsy (Williams, 1953; Babb and Crandall, 1976; Valentin et al., 2002; Valentin et al., 2005). The limbic H wave morphology as presented in Figure 3 resembles to some extend an interictal spike-wave discharge, which is commonly seen in both scalp and invasive EEG monitoring. Spontaneous examples may occur as single examples or in trains. The observed limbic H-wave contains a single cycle, had reliable latency across trials and subjects and shows specificity to the PCC. While the epilepsy and a hyperexcitable limbic system may therefore facilitate this response, future research will have to determine to what extent it is only observed in patients with epilepsy.

## Conclusion

Single pulse electrical stimulation reveals the limbic H-wave in the PCC after stimulating the hippocampus. Combined with diffusion imaging, and well established anatomical knowledge (Papez, 1937; Wang et al., 2020), it is likely that this response is generated by indirect projections through the hippocampal limbic system. Stimulation-generated waveforms have been used for many decades as electrophysiological biomarkers of motor system function. These data suggest that the limbic H-wave we describe can be used as such an electrophysiological biomarker of the memory-spatial part of the limbic system.

## Acknowledgements

The research project was supported by the Mayo Clinic DERIVE Office and Mayo Clinic Center for Biomedical Discovery support and the National Institute Of Mental Health of the National Institutes of Health under Award Number R01MH122258. The content is solely the responsibility of the authors and does not necessarily represent the official views of the National Institutes of Health. We are very grateful to all the subjects enrolled in the study, and the assistance provided by Cindy Nelson and Karla Crocket is greatly appreciated.

## Glossary

PCC: Posterior cingulate cortex
SPES: Single pulse electrical stimulation
CCEP: Cortico-cortical evoked potential
ANT: Anterior nucleus of the thalamus
MEG: Magnetoencephalography
EEG: Electroencephalography
sEEG: Stereo electroencephalography
SOZ: Seizure Onset Zone
Hc: Hippocampus
PH: Parahippocampal gyrus
HC: Hippocampal Complex
Amg: Amygdala
ACC: Anterior cingulate cortex
MAC: Middle-anterior cingulate cortex
MPC: Middle-posterior cingulate cortex
PDC: Post-dorsal cingulate cortex
PVC: Post-ventral cingulate cortex
Tha: Thalamus
PCA: Principal component analysis.

## Supplementary Figures

**Figure 2-1.**
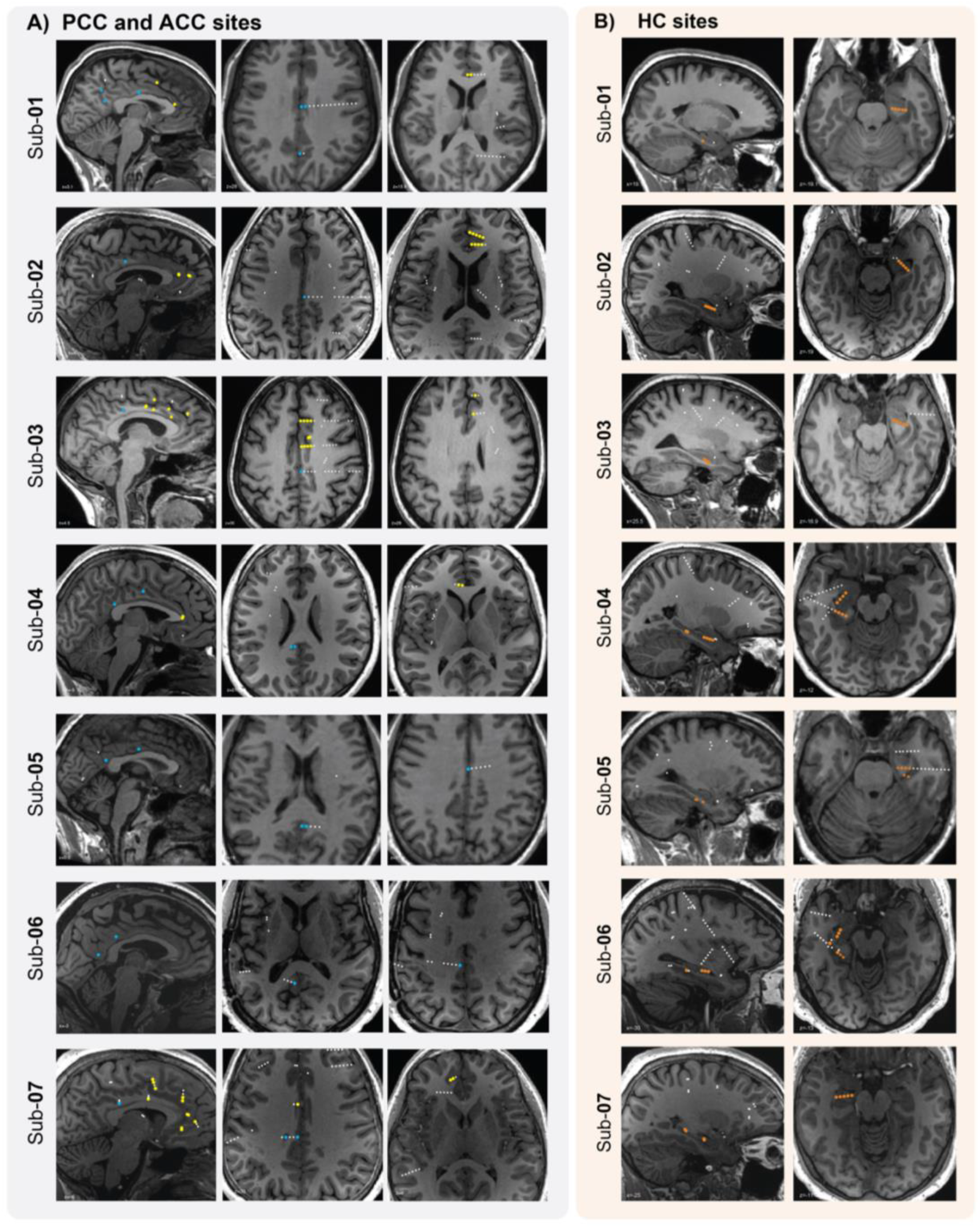
Anatomical estimation of cingulate cortex and HC electrode contacts in T1w brain slices across subjects. **A)** Sagittal (left panel) and axial (right panel) view of the electrode estimations implanted in the HC (in orange, circles for hippocampus and stars for parahippocampal gyrus), **B)** Sagittal (left panel) and axial (right panel) view of the electrode estimations implanted in the amygdala (purple circles), **C)** Sagittal (left panel) and axial (right panel) view of the electrode estimations implanted in the ANT (green circles).

**Figure 2-2.**
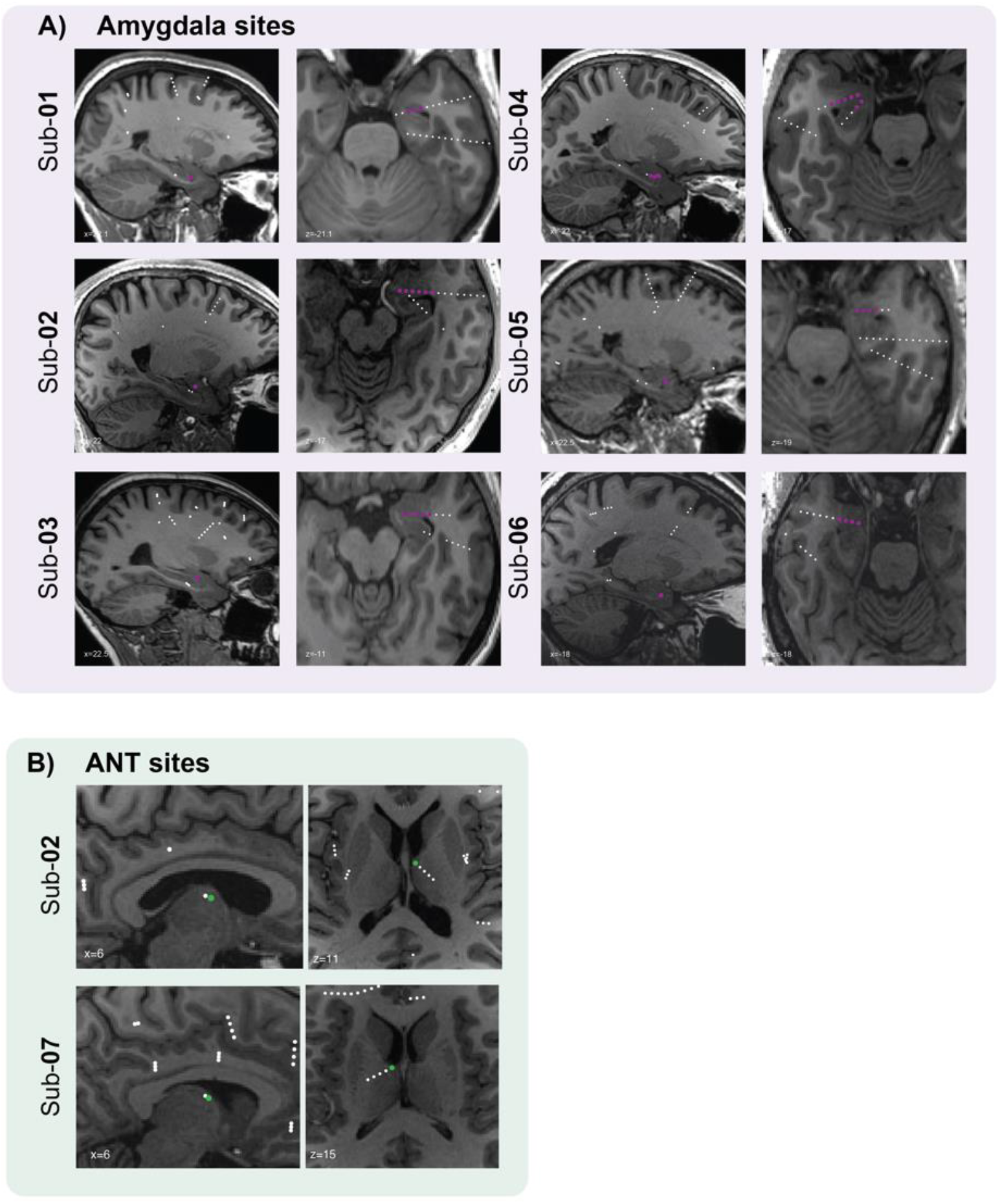
Anatomical estimation of amygdala and ANT electrode contacts in T1w brain slices across subjects. **A)** Sagittal (left panel) and axial (right panel) view of the electrode estimations implanted in the amygdala, and **B)** in the ANT.

**Figure 2-3.**
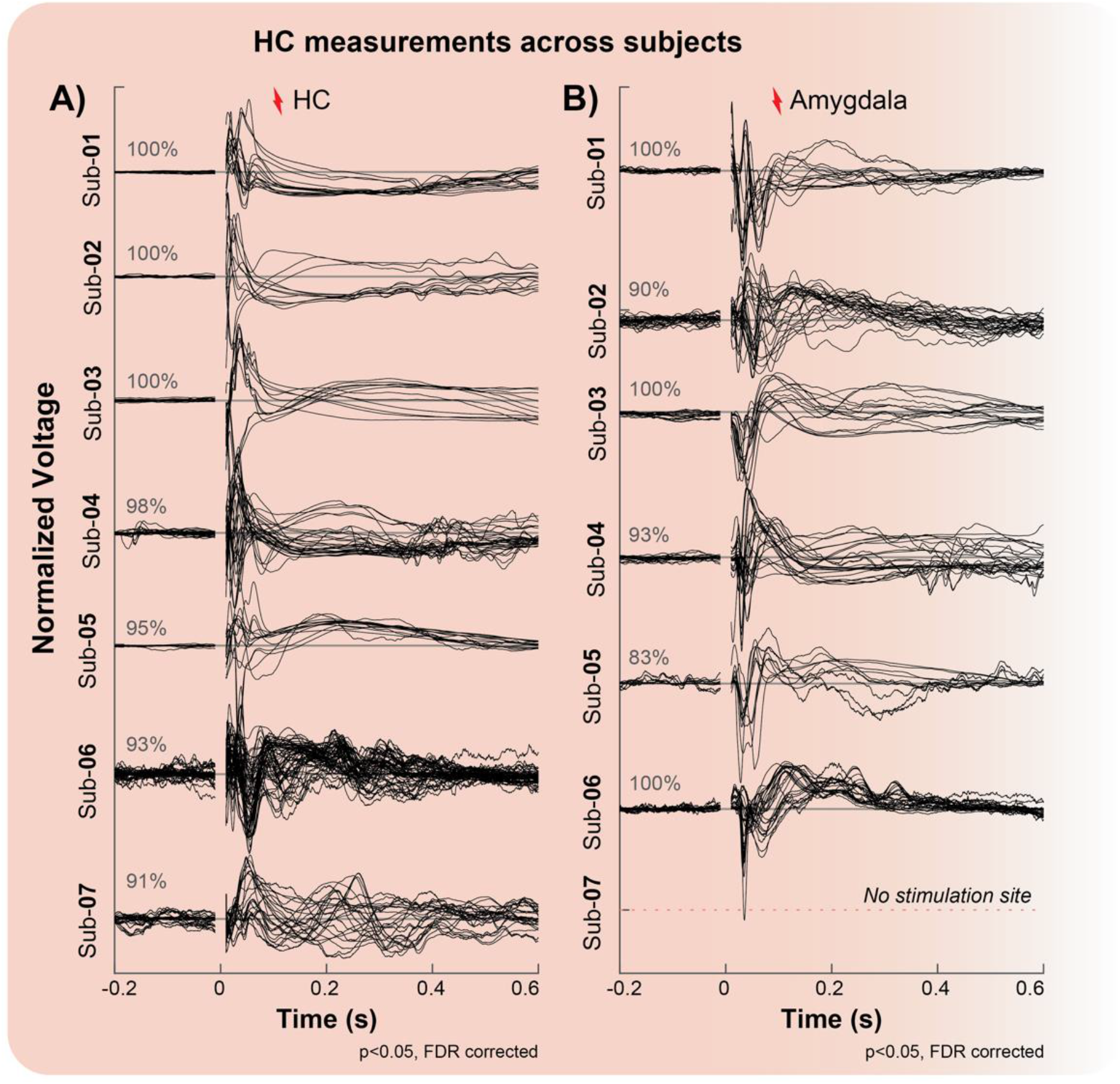
Measurements in the HC. Significant CCEPs in the HC (t-tests, p_FDRcorrected_<0.05, L2-normalized) are represented by each black line after stimulation in two different regions denoted in A) and B). **A)** Recordings in the HC after stimulation within the HC with a mean of 97% of significant CCEPs across subjects. **B)** Recordings in the HC upon stimulation in the with a mean of 94% of significant CCEPs across subjects.

**Figure 2-4.**
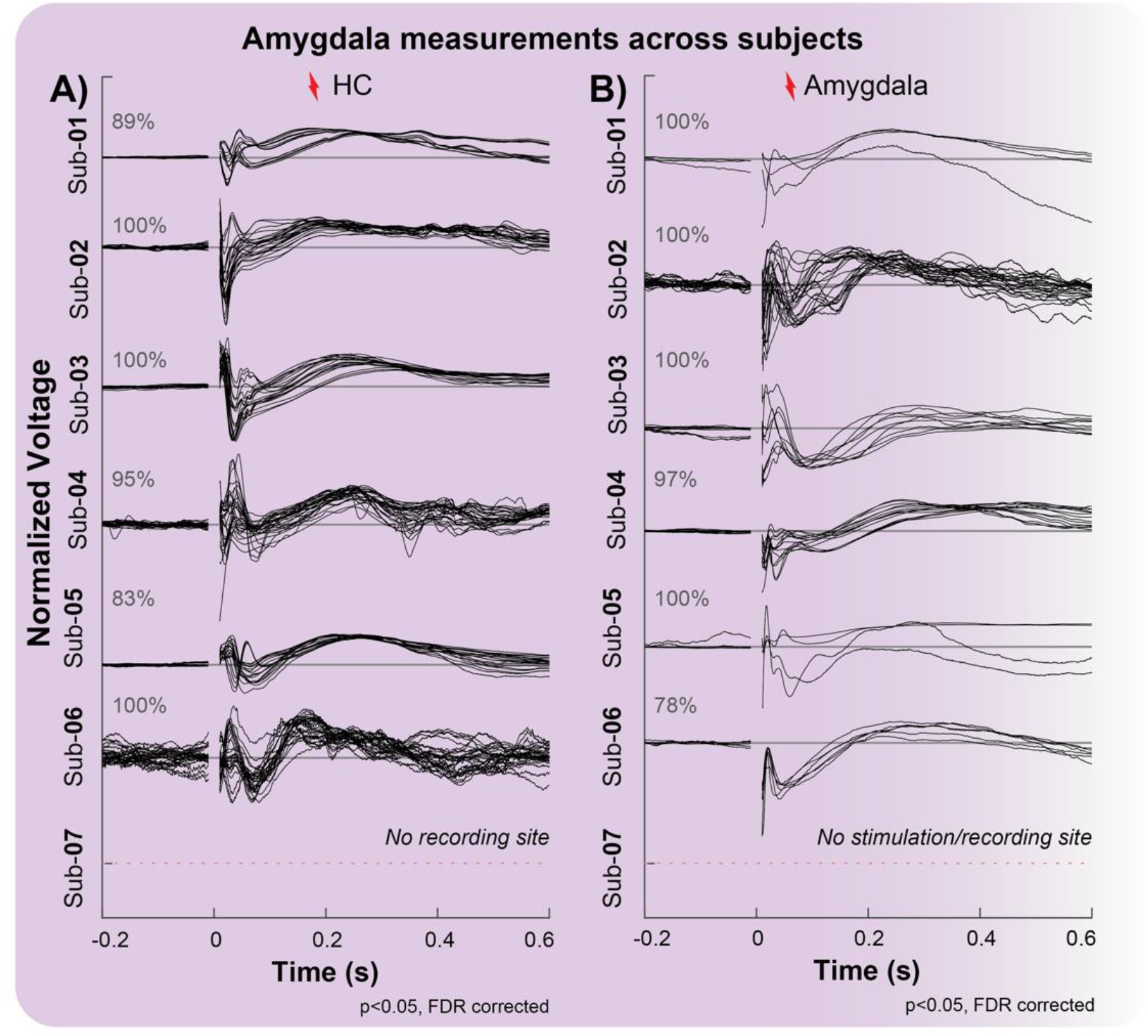
Measurements in the amygdala. Significant CCEPs in the amygdala (t-tests, p_FDRcorrected_<0.05, L2-normalized) are represented by each black line after stimulation in two different regions denoted in A) and B). **A)** Recordings in amygdala upon stimulation in the HC with a mean of 95% of significant CCEPs across subjects. **B)** Recordings in amygdala after stimulation within the amygdala with a mean of 96% of significant CCEPs across subjects.

**Figure 4-1.**
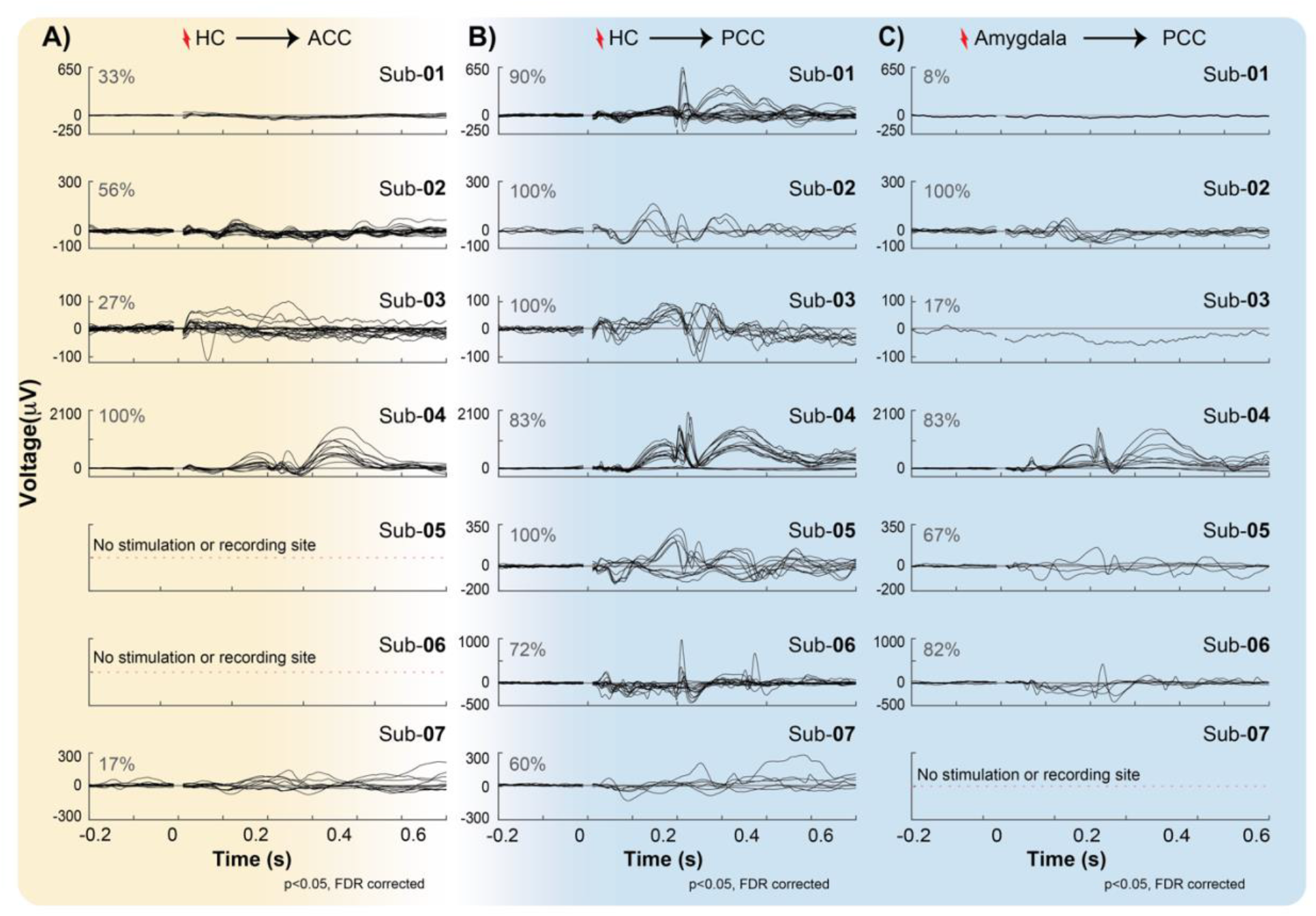
CCEPs measured from PCC and ACC after hippocampal and amygdala stimulation with voltage in μV. Significant average responses (t-tests, pFDRcorrected<0.05) are represented by each black line under three different conditions, denoted inA), B), and C). **A)** ACC under hippocampal stimulation, **B)** PCC under hippocampal stimulation, and **C)** PCC under amygdala stimulation.

## References

(1998) History of Neuroscience in Autobiography Vol. 2. 525 B Street, Suite 1900, San Diego, California 92101-4495, USA: The Society for Neuroscience.

Aggleton JP, Nelson AJD, O’Mara SM (2022) Time to retire the serial Papez circuit: Implications for space, memory, and attention. Neurosci Biobehav Rev 140:104813.

Aponik-Gremillion L, Chen YY, Bartoli E, Koslov SR, Rey HG, Weiner KS, Yoshor D, Hayden BY, Sheth SA, Foster BL (2022) Distinct population and single-neuron selectivity for executive and episodic processing in human dorsal posterior cingulate. Elife 11.

Awiszus F, Feistner H (1994) Quantification of D-and I-wave effects evoked by transcranial magnetic brain stimulation on the tibialis anterior motoneuron pool in man. Experimental Brain Research 101:153–158.

Babb TL, Crandall PH (1976) Epileptogenesis of human limbic neurons in psychomotor epileptics. Electroencephalogr Clin Neurophysiol 40:225–243.

Beckmann M, Johansen-Berg H, Rushworth MF (2009) Connectivity-based parcellation of human cingulate cortex and its relation to functional specialization. J Neurosci 29:1175–1190.

Bertram EH, Zhang DX, Mangan P, Fountain N, Rempe D (1998) Functional anatomy of limbic epilepsy: a proposal for central synchronization of a diffusely hyperexcitable network. Epilepsy Res 32:194–205.

Borchers S, Himmelbach M, Logothetis N, Karnath H-O (2012) Direct electrical stimulation of human cortex, gold standard for mapping brain functions. Nature Reviews Neuroscience 13:63–70.

Boyd SG, Rothwell JC, Cowan JMA, Webb PJ, Morley T, Asselman P, Marsden CD (1986) A Method of Monitoring Function in Corticospinal Pathways during Scoliosis Surgery with a Note on Motor Conduction Velocities. J Neurol Neurosur Ps 49:251–257.

Brunton SL, Kutz JN (2022) Data-driven science and engineering: Machine learning, dynamical systems, and control: Cambridge University Press.

Bubb EJ, Kinnavane L, Aggleton JP (2017) Hippocampal - diencephalic - cingulate networks for memory and emotion: An anatomical guide. Brain Neurosci Adv 1.

Bubb EJ, Metzler-Baddeley C, Aggleton JP (2018) The cingulum bundle: Anatomy, function, and dysfunction. Neurosci Biobehav Rev 92:104–127.

Carmichael ST, Price JL (1996) Connectional networks within the orbital and medial prefrontal cortex of macaque monkeys. J Comp Neurol 371:179–207.

Child ND, Benarroch EE (2013) Anterior nucleus of the thalamus: functional organization and clinical implications. Neurology 81:1869–1876.

Cieslak M et al. (2021) QSIPrep: an integrative platform for preprocessing and reconstructing diffusion MRI data. Nat Methods 18:775–778.

Costa P, Peretta P, Faccani G (2013) Relevance of intraoperative D wave in spine and spinal cord surgeries. Eur Spine J 22:840–848.

Daubechies I (1992) Ten lectures on wavelets. Philadelphia, Pennsylvania

Destrieux C, Fischl B, Dale A, Halgren E (2010) Automatic parcellation of human cortical gyri and sulci using standard anatomical nomenclature. Neuroimage 53:1–15.

Donos C, Maliia MD, Mindruta I, Popa I, Ene M, Balanescu B, Ciurea A, Barborica A (2016) A connectomics approach combining structural and effective connectivity assessed by intracranial electrical stimulation. Neuroimage 132:344–358.

Efron B, Tibshirani R (1998) An introduction to the bootstrap. Boca Raton; London: Chapman & Hall/CRC.

Enatsu R, Gonzalez-Martinez J, Bulacio J, Kubota Y, Mosher J, Burgess RC, Najm I, Nair DR (2015) Connections of the Limbic Neetwork: A corticocortical evoked potentials study. Cortex 62:20–33.

Fischl B (2012) FreeSurfer. Neuroimage 62:774–781.

Foo TKF et al. (2018) Lightweight, compact, and high-performance 3T MR system for imaging the brain and extremities. Magn Reson Med 80:2232–2245.

Friston KJ, Ashburner J, Kiebel S, Nichols T, Penny WD, ScienceDirect (2007) Statistical parametric mapping: the analysis of funtional brain images, First edition. Edition. Amsterdam; Boston: Elsevier/Academic Press.

Gregg NM, Sladky V, Nejedly P, Mivalt F, Kim I, Balzekas I, Sturges BK, Crowe C, Patterson EE, Van Gompel JJ, Lundstrom BN, Leyde K, Denison TJ, Brinkmann BH, Kremen V, Worrell GA (2021) Thalamic deep brain stimulation modulates cycles of seizure risk in epilepsy. Sci Rep 11:24250.

Grewal SS, Middlebrooks EH, Kaufmann TJ, Stead M, Lundstrom BN, Worrell GA, Lin C, Baydin S, Van Gompel JJ (2018) Fast gray matter acquisition T1 inversion recovery MRI to delineate the mammillothalamic tract for preoperative direct targeting of the anterior nucleus of the thalamus for deep brain stimulation in epilepsy. Neurosurg Focus 45.

Gronlier E, Vendramini E, Volle J, Wozniak-Kwasniewska A, Anton Santos N, Coizet V, Duveau V, David O (2021) Single-pulse electrical stimulation methodology in freely moving rat. J Neurosci Methods 353:109092.

Gupta MR, Jacobson NP (2006) Wavelet principal component analysis and its application to hyperspectral images. In: 2006 International Conference on Image Processing, pp 1585 – 1588. Atlanta, GA.

Hermes D, Miller KJ, Noordmans HJ, Vansteensel MJ, Ramsey NF (2010) Automated electrocorticographic electrode localization on individually rendered brain surfaces. J Neurosci Methods 185:293–298.

Hicks RG, Burke DJ, Stephen JPH (1991) Monitoring Spinal-Cord Function during Scoliosis Surgery with Cotrel-Dubousset Instrumentation. Med J Australia 154:82–86.

Holtzheimer PE, Kelley ME, Gross RE, Filkowski MM, Garlow SJ, Barrocas A, Wint D, Craighead MC, Kozarsky J, Chismar R, Moreines JL, Mewes K, Posse PR, Gutman DA, Mayberg HS (2012) Subcallosal cingulate deep brain stimulation for treatment-resistant unipolar and bipolar depression. Arch Gen Psychiatry 69:150–158.

In MH, Tan ET, Trzasko JD, Shu Y, Kang D, Yarach U, Tao S, Gray EM, Huston J, 3rd, Bernstein MA (2020) Distortion-free imaging: A double encoding method (DIADEM) combined with multiband imaging for rapid distortion-free high-resolution diffusion imaging on a compact 3T with high-performance gradients. J Magn Reson Imaging 51:296–310.

Innocenti GM, Vercelli A, Caminiti R (2014) The diameter of cortical axons depends both on the area of origin and target. Cereb Cortex 24:2178–2188.

Jo HJ, Kenney-Jung DL, Balzekas I, Welker KM, Jones DT, Croarkin PE, Benarroch EE, Worrell GA (2019) Relationship Between Seizure Frequency and Functional Abnormalities in Limbic Network of Medial Temporal Lobe Epilepsy. Front Neurol 10:488.

Jones DK, Christiansen KF, Chapman RJ, Aggleton JP (2013) Distinct subdivisions of the cingulum bundle revealed by diffusion MRI fibre tracking: implications for neuropsychological investigations. Neuropsychologia 51:67–78.

Kahn L, Sutton B, Winston HR, Abosch A, Thompson JA, Davis RA (2021) Deep Brain Stimulation for Obsessive-Compulsive Disorder: Real World Experience Post-FDA-Humanitarian Use Device Approval. Front Psychiatry 12:568932.

Keller CJ, Honey CJ, Megevand P, Entz L, Ulbert I, Mehta AD (2014) Mapping human brain networks with cortico-cortical evoked potentials. Philos Trans R Soc Lond B Biol Sci 369.

Koubeissi MZ, Kahriman E, Syed TU, Miller J, Durand DM (2013) Low-frequency electrical stimulation of a fiber tract in temporal lobe epilepsy. Annals of Neurology 74:223–231.

Kubota Y, Enatsu R, Gonzalez-Martinez J, Bulacio J, Mosher J, Burgess RC, Nair DR (2013) In vivo human hippocampal cingulate connectivity: a corticocortical evoked potentials (CCEPs) study. Clin Neurophysiol 124:1547–1556.

Kumaravelu K, Oza CS, Behrend CE, Grill WM (2018) Model-based deconstruction of cortical evoked potentials generated by subthalamic nucleus deep brain stimulation. Journal of Neurophysiology 120:662–680.

Lockman J, Fisher RS (2009) Therapeutic brain stimulation for epilepsy. Neurol Clin 27:1031–1040.

Lozano AM, Lipsman N, Bergman H, Brown P, Chabardes S, Chang JW, Matthews K, McIntyre CC, Schlaepfer TE, Schulder M, Temel Y, Volkmann J, Krauss JK (2019) Deep brain stimulation: current challenges and future directions. Nat Rev Neurol 15:148–160.

Luo Y, Sun Y, Tian X, Zheng X, Wang X, Li W, Wu X, Shu B, Hou W (2021) Deep Brain Stimulation for Alzheimer’s Disease: Stimulation Parameters and Potential Mechanisms of Action. Front Aging Neurosci 13:619543.

Matsumoto R, Nair DR, LaPresto E, Najm I, Bingaman W, Shibasaki H, Luders HO (2004) Functional connectivity in the human language system: a cortico-cortical evoked potential study. Brain 127:2316–2330.

Miller KJ, Muller KR, Hermes D (2021) Basis profile curve identification to understand electrical stimulation effects in human brain networks. PLoS Comput Biol 17:e1008710.

Miller KJ, Prieto T, Williams NR, Halpern CH (2019) Case Studies in Neuroscience: The electrophysiology of a human obsession in nucleus accumbens. J Neurophysiol 121:2336–2340.

Mitzdorf U (1985) Current Source-Density Method and Application in Cat Cerebral-Cortex - Investigation of Evoked-Potentials and Eeg Phenomena. Physiol Rev 65:37–100.

Mufson EJ, Pandya DN (1984) Some Observations on the Course and Composition of the Cingulum Bundle in the Rhesus-Monkey. J Comp Neurol 225:31–43.

Nair DR et al. (2020) Nine-year prospective efficacy and safety of brain-responsive neurostimulation for focal epilepsy. Neurology 95:e1244–e1256.

Nemanic S, Alvarado MC, Bachevalier J (2004) The hippocampal/parahippocampal regions and recognition memory: Insights from visual paired comparison versus object-delayed nonmatching in monkeys. Journal of Neuroscience 24:2013–2026.

Oane I, Barborica A, Chetan F, Donos C, Maliia MD, Arbune AA, Daneasa A, Pistol C, Nica AE, Bajenaru OA, Mindruta I (2020) Cingulate cortex function and multi-modal connectivity mapped using intracranial stimulation. Neuroimage 220:117059.

Pal Attia T, Crepeau D, Kremen V, Nasseri M, Guragain H, Steele SW, Sladky V, Nejedly P, Mivalt F, Herron JA, Stead M, Denison T, Worrell GA, Brinkmann BH (2021) Epilepsy Personal Assistant Device-A Mobile Platform for Brain State, Dense Behavioral and Physiology Tracking and Controlling Adaptive Stimulation. Front Neurol 12:704170.

Papez JW (1937) A Proposed Mechanism of Emotion. Archieves of Neurology and Psychiatry 38:725–774.

Patton HD, Amassian VE (1954) Single and multiple-unit analysis of cortical stage of pyramidal tract activation. J Neurophysiol 17:345–363.

Pearce PS, Friedman D, Lafrancois JJ, Iyengar SS, Fenton AA, Maclusky NJ, Scharfman HE (2014) Spike-wave discharges in adult Sprague-Dawley rats and their implications for animal models of temporal lobe epilepsy. Epilepsy Behav 32:121–131.

Puyati W, Walairacht S, Walairacht A (2006) PCA in wavelet domain for face recognition. In: 2006 8th International Conference Advanced Communication Technology. Boca Raton Phoenix Park, Korea (South).

Quinones-Hinojosa A, Lyon R, Zada G, Lamborn KR, Gupta N, Parsa AT, McDermott MW, Weinstein PR (2005) Changes in transcranial motor evoked potentials during intramedullary spinal cord tumor resection correlate with postoperative motor function. Neurosurgery 56:982–993; discussion 982-993.

Rolls ET (2015) Limbic systems for emotion and for memory, but no single limbic system. Cortex 62:119–157.

Salanova V et al. (2015) Long-term efficacy and safety of thalamic stimulation for drug-resistant partial epilepsy. Neurology 84:1017–1025.

Siddiqi SH et al. (2021) Brain stimulation and brain lesions converge on common causal circuits in neuropsychiatric disease. Nat Hum Behav 5:1707–1716.

Strange BA, Witter MP, Lein ES, Moser EI (2014) Functional organization of the hippocampal longitudinal axis. Nat Rev Neurosci 15:655–669.

Takeyama H, Matsumoto R, Usami K, Nakae T, Kobayashi K, Shimotake A, Kikuchi T, Yoshida K, Kunieda T, Miyamoto S, Takahashi R, Ikeda A (2019) Human entorhinal cortex electrical stimulation evoked short-latency potentials in the broad neocortical regions: Evidence from cortico-cortical evoked potential recordings. Brain Behav 9:e01366.

Valentin A, Anderson M, Alarcon G, Seoane JJ, Selway R, Binnie CD, Polkey CE (2002) Responses to single pulse electrical stimulation identify epileptogenesis in the human brain in vivo. Brain 125:1709–1718.

Valentin A, Alarcon G, Honavar M, Garcia Seoane JJ, Selway RP, Polkey CE, Binnie CD (2005) Single pulse electrical stimulation for identification of structural abnormalities and prediction of seizure outcome after epilepsy surgery: a prospective study. Lancet Neurol 4:718–726.

Vogt BA (2019) Cingulate cortex in the three limbic subsystems. Handb Clin Neurol 166:39–51.

Wang YC, Kremen V, Brinkmann BH, Middlebrooks EH, Lundstrom BN, Grewal SS, Guragain H, Wu MH, Van Gompel JJ, Klassen BT, Stead M, Worrell GA (2020) Probing circuit of Papez with stimulation of anterior nucleus of the thalamus and hippocampal evoked potentials. Epilepsy Res 159:106248.

Willbrand EH, Parker BJ, Voorhies WI, Miller JA, Lyu I, Hallock T, Aponik-Gremillion L, Koslov SR, Alzheimer’s Disease Neuroimaging I, Bunge SA, Foster BL, Weiner KS (2022) Uncovering a tripartite landmark in posterior cingulate cortex. Sci Adv 8:eabn9516.

Williams D (1953) A study of thalamic and cortical rhythms in petit mal. Brain 76:50–69.

Wu H, Miller KJ, Blumenfeld Z, Williams NR, Ravikumar VK, Lee KE, Kakusa B, Sacchet MD, Wintermark M, Christoffel DJ, Rutt BK, Bronte-Stewart H, Knutson B, Malenka RC, Halpern CH (2018) Closing the loop on impulsivity via nucleus accumbens delta-band activity in mice and man. Proceedings of the National Academy of Sciences 115:192–197.

Wu Y, Sun D, Wang Y, Wang Y, Ou S (2016) Segmentation of the Cingulum Bundle in the Human Brain: A New Perspective Based on DSI Tractography and Fiber Dissection Study. Front Neuroanat 10:84.

Wyllie E (2012) Wyllie’s treatment of epilepsy: Principles and practice: Fifth edition.

Yeh FC (2020) Shape analysis of the human association pathways. Neuroimage 223:117329.

Yeh FC, Wedeen VJ, Tseng WY (2010) Generalized q-sampling imaging. IEEE Trans Med Imaging 29:1626–1635.

Yeh FC, Liu L, Hitchens TK, Wu YL (2017) Mapping immune cell infiltration using restricted diffusion MRI. Magn Reson Med 77:603–612.

Yeh FC, Verstynen TD, Wang Y, Fernandez-Miranda JC, Tseng WY (2013) Deterministic diffusion fiber tracking improved by quantitative anisotropy. PLoS One 8:e80713.

Yeh FC, Panesar S, Barrios J, Fernandes D, Abhinav K, Meola A, Fernandez-Miranda JC (2019) Automatic Removal of False Connections in Diffusion MRI Tractography Using Topology-Informed Pruning (TIP). Neurotherapeutics 16:52–58.

Yeh FC, Panesar S, Fernandes D, Meola A, Yoshino M, Fernandez-Miranda JC, Vettel JM, Verstynen T (2018) Population-averaged atlas of the macroscale human structural connectome and its network topology. Neuroimage 178:57–68.

